# The atypical antidepressant tianeptine causes opioid-receptor-dependent beta oscillations in the rat hippocampus

**DOI:** 10.64898/2025.11.30.691361

**Authors:** Scott G Burt, Grant D Phillips, Jeremy J Lambert, Stephen J Martin

## Introduction

According to the 2021 Global Burden of Diseases study, major depressive disorder (MDD) is currently the second largest worldwide contributor to years lived with disability (Ferrari, Santomauro, et al., 2024), and its prevalence is growing rapidly, particularly among adolescents and young adults (Zhang et al., 2025). Current first-line pharmacological treatment typically involves selective serotonin reuptake inhibitors (SSRIs), but these drugs have variable efficacy, therapeutic benefits can take weeks to arise, and cessation of treatment can result in a distressing ‘discontinuation syndrome’ (Sharp and Collins, 2024). To meet the clinical need for fast-acting and more effective treatments, several drugs with atypical mechanisms of action have recently entered clinical use or evaluation. These include ketamine (Berman et al., 2000; Zanos & Gould 2018), psychedelics such as psilocybin (Wulff et al., 2023; Nutt et al., 2024), and neurosteroids such as allopregnanolone and related compounds (Belelli et al., 2022; Ahmad et al., 2024).

Another antidepressant drug with an atypical mechanism of action and relatively rapid therapeutic effects is tianeptine (Novotny & Faltus, 2002; Han et al., 2022). Although not a novel compound—it was first licensed for clinical use in France in the 1980s (Nishio et al., 2024)—its molecular mechanism of action was poorly understood until just over a decade ago when it was identified as an agonist of µ and δ-opioid receptors (µ-ORs and δ-ORs; Gassaway et al., 2014). This was unexpected, but not without precedent. Until the mid-20^th^ century, opium was recognised as a fast and effective treatment for ‘melancholia’ or melancholic depression (Jelen et al., 2022), now viewed as a severe sub-type of MDD, often resistant to treatment with SSRIs (e.g. Parker, 2017). More recently, evidence has emerged of the antidepressant efficacy of other opioids such as buprenorphine (Fava et al., 2016; Robinson et al., 2017; Bhivandkar et al., 2024), and of selective δ-opioid receptor agonists (Jelen et al., 2024).

In addition to its antidepressant actions, tianeptine also improves memory in both rodents and humans (Morris et al., 2001, Zoladz et al., 2008; McEwen et al., 2010; Jeon et al., 2014), and a recent retrospective analysis reveals its efficacy in improving cognitive function in Alzheimer’s Disease (Garcia-Alberca et al., 2022). Many studies have identified neurotrophic effects following tianeptine administration, such as increased BDNF expression (Zoladz et al., 2008), increased dendritic growth and excitatory synapse formation (McEwen et al., 2002; Chu et al., 2010), and the prevention of stress-induced deficits in synaptic plasticity (Dupin et al., 2006; Zoladz et al., 2008) and hippocampal volume (Pollano et al., 2016). Tianeptine also enhances hippocampal AMPA-receptor-mediated synaptic transmission, stabilises the surface diffusion of synaptic AMPA receptors, and facilitates the induction of hippocampal long-term potentiation (LTP) of synaptic strength (Kole et al., 2002; Szegedi et al., 2011; Pillai et al., 2012; Zhang et al., 2013 & 2018). Consistent with these actions, tianeptine increases the phosphorylation of AMPA receptor GluA1 subunits at serine 831 and 845, sites for the action of Ca^2+/^calmodulin-dependent protein kinase II (CaMKII) and protein kinase A (PKA) respectively (Svenningsson et al., 2007; Szegedi et al., 2011). Since the discovery of its opioid receptor agonist properties, µ-opioid receptors on somatostatin (STT^+^) interneurons of the hippocampus have been identified as key targets underlying tianeptine’s antidepressant effects (Han et al., 2022), underscoring the therapeutic potential of opioid receptors as novel antidepressant targets. Since the activation of µ-opioid receptors on inhibitory interneurons is expected to result in a disinhibition of principal neurons, it is possible that tianeptine exerts dual effects on hippocampal excitatory transmission by simultaneously enhancing the activation of AMPA receptors on principal neurons and reducing GABA release from interneurons. Both phenomena might be relevant to tianeptine’s facilitation of LTP, but effects on oscillatory network activity are also likely owing to the role of mutually interconnected networks of inhibitory interneurons and excitatory neurons in the generation of hippocampal oscillations such as gamma rhythms (Colgin & Moser, 2010; Antonoudiou et al., 2020).

To address this last possibility, we investigated the effects of tianeptine administration on local field potential (LFP) activity in area CA1 of intact rats. Since tianeptine enhances hippocampal fast synaptic transmission in hippocampal slices, we also recorded fEPSPs evoked in CA1 by stimulation of CA3 to assess the effects of the drug in vivo. For comparison, we studied the effects of buprenorphine, another opioid receptor agonist with antidepressant actions (Emrich et al., 1982; Bhivandkar et al., 2024), and of ketamine, a fast-acting atypical antidepressant known to enhance hippocampal gamma oscillations (Ma & Leung, 2007; Caixeta et al., 2013). To determine the role of opioid receptors in any changes observed, we tested the effects of co-administration of each drug with the opioid receptor antagonist naloxone. To control for possible drug-induced changes in brain temperature, we implanted a miniature temperature probe in the hippocampus contralateral to the stimulating and recording electrodes in a subset of animals.

## Materials and Methods

### Animals

All procedures involving animals were conducted in accordance with the UK Animals (Scientific Procedures) Act (1986) and subject to local ethical review by the University of Dundee’s Welfare and Ethical Use of Animals in Research Committee. Male Lister hooded rats obtained from Charles River UK were used in all experiments. Prior to the experiment, rats were housed in groups of 4 in pairs of cages, each 32 x 50 cm in area, and connected by an acrylic tube. They were given unrestricted access to food (standard raw chow, sunflower seeds, and wheat grains) and water, and maintained on a 12-h light / 12-h dark cycle at constant temperature (19-24°C). Bedding and nesting materials comprised wood shavings, paper wool, ‘sizzle-nest’, and hay, and enrichment items included acrylic tubes for refuge, wooden chew-sticks, and Aspen balls,

### Drugs and their administration

All drugs were dissolved in sterile injectable saline (hereafter referred to as ‘vehicle’) and injected subcutaneously at a volume of 5ml/kg. The drugs and doses used were as follows: tianeptine (10 mg/kg & 30 mg/kg; Kemprotec Limited, Cumbria, UK), (±)-ketamine hydrochloride (10 mg/kg; Sigma-Aldrich, Burlington, MA, USA), buprenorphine (0.03 mg/kg; Sigma-Aldrich), and naloxone (1 mg/kg; Sigma-Aldrich).

### Surgery

At the start of each experiment, the rat was anaesthetised with 1.5 g/kg urethane (ethyl carbamate; 0.3 mg/ml; 5 ml/kg; IP) and placed in a stereotaxic frame (Kopf, Tujunga, USA) with the skull horizontal. Body temperature was monitored via a rectal probe and maintained at 36.2°C using a homeothermic heat blanket (Harvard Apparatus, Holliston, MA, USA). Depth of anaesthesia was assessed throughout the experiment, and 0.2 ml top-up doses of urethane were administered as required. Small holes were drilled in the skull for the insertion of the electrodes and temperature probe. A small stainless steel jeweller’s screw was secured to the occipital bone to serve as a reference for fEPSP and LFP recording.

### Recording of field excitatory postsynaptic potentials (fEPSPs)

A PTFE-insulated monopolar platinum/iridium recording electrode (external diameter = 0.103 mm) was lowered into the left hippocampus targeting the stratum radiatum of area CA1 (3.8 mm posterior and 2.5 mm lateral to bregma; depth circa -2.4 mm from the dura). A bipolar stimulating electrode comprising two twisted wires identical in composition to the recording electrode was lowered into ipsilateral area CA3 (3.5 mm posterior and 3.0 mm lateral to bregma; depth circa -3.0 mm from the dura) to activate the Schaffer collateral projection to CA1. A thermistor (external diameter = 0.4 mm) was lowered into the right hippocampus at the same coordinates as the recording electrode in the left hippocampus. Fig. 1 shows a schematic illustration of the electrode locations. Correct electrode placement was verified by the characteristic depth profile of fEPSP changes during insertion (see Shires et al., 2012, Fig. 6 for a detailed explanation).

**Fig. 1.**
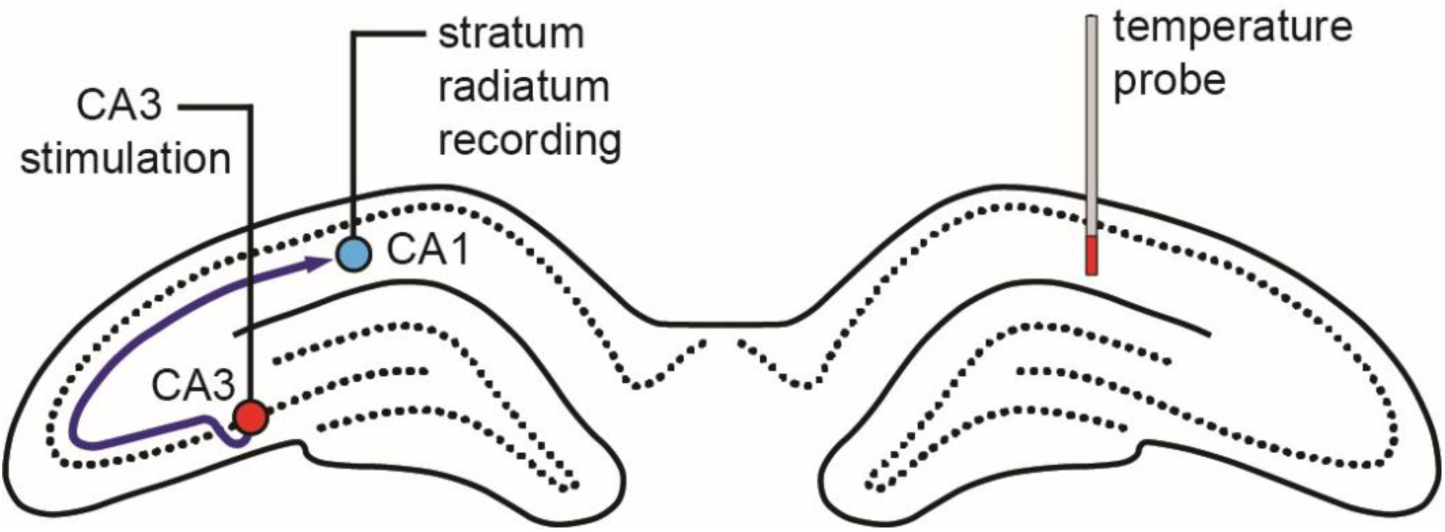
Schematic illustration of the placement of stimulating and recording electrodes in CA3 and CA1 stratum radiatum respectively, and a temperature probe in the contralateral CA1 stratum radiatum.

Evoked field excitatory postsynaptic potentials (fEPSPs) were amplified and filtered (high pass = 0.3 Hz; low pass = 5 kHz) using a differential AC amplifier (model 3500, A-M Systems, Sequim, WA, USA) and sampled at 20 kHz using a data acquisition card (PCIe-6321; National Instruments, Austin, TX, USA) mounted in a PC running custom-written LabView software for the control of electrical stimulation and the time-locked recording of evoked fEPSPs (Evoked Potential Sampler, Patrick Spooner, University of Edinburgh). 50-Hz line noise was removed using a HumBug noise-elimination unit (Digitimer, Welwyn Garden City, UK). Stimulation was delivered via a Neurolog system and stimulus isolator units (Digitimer), and consisted of biphasic constant-current pulses, 100 µs per phase. At the start of each experiment, electrodes were lowered into the hippocampus, and depths were adjusted to maximise the amplitude of the negative-going dendritic fEPSP elicited in the stratum radiatum of CA1 by stimulation of CA3 (see Shires et al., 2012). Stimulation intensity was adjusted to elicit a fEPSP with a slope of ∼50% of the maximum value that could be obtained during an initial input-output curve; stimulation currents fell within the range of 200-500 µA in all cases.

After electrode placement, baseline recording comprising single test pulses delivered at 20-s intervals began. Once stable fEPSPs had been recorded for at least 30 min, an input-output curve was conducted, and baseline recording continued for another 20 min before drug injection. All drugs were injected subcutaneously (SC) via a butterfly needle secured in place at the start of the experiment to avoid the need for needle insertion at the time of injection. 30 min after injection, a second input-output curve was carried out, followed by a further 30-min period of recording.

For each fEPSP, its slope was calculated online by the acquisition software, measured by linear regression between two fixed time-points corresponding to the early rising phase of the waveform. The amplitude of the tail portion of each fEPSP was measured offline by determining the peak positive amplitude of the ‘overshoot’ portion of the waveform, a measure that reflects the recruitment of feedforward inhibition (Fig. S1; cf. Alger & Nicoll, 1982; Arai et al., 1995; Karnup & Stelzer, 1999). fEPSP slope and tail amplitude measures were normalised to the mean value over the 10-min period before drug injection (assigned a value of 100%), and mean values were calculated over successive 2-min recording periods for the remainder of the experiment.

### Spontaneous local field potential (LFP) recording

In all experiments, the continuous wide-band LFP was recorded in parallel to the fEPSPs, via the same recording electrode, by splitting the output of the differential amplifier and sampling the data using a USB-6003 acquisition device (National Instruments, Austin, TX, USA) connected to a PC running custom-written software for LFP capture (Patrick Spooner, University of Edinburgh). Data were acquired at 20 kHz, low-pass filtered at 100 Hz and down-sampled to 200 Hz. Successive 2-s samples of data (50% window overlap; temporally filtered using the Hann function) were analysed by fast Fourier transform (FFT). Samples containing fEPSPs were omitted from the analysis. Mean power spectral density from 1-80 Hz was analysed in 0.5 Hz frequency bins, and averaged across successive 2-min recording periods. Data were converted to log_10_ values. Time-frequency plots were constructed by plotting successive 2-min power spectra as a 3-D heat map with power as the z-axis. The change in spectral power before and after drug injection was calculated by subtracting the log_10_ power at each time point and frequency bin from the corresponding mean over the 10-min period before injection. These data were also plotted as a 3-D time-frequency graph, this time with change in spectral power relative to baseline as the z-axis. The mean change in power was also calculated in specific frequency bands—theta (3-4 Hz), beta (10-30 Hz), slow gamma (30-50 Hz) and fast gamma (50-80 Hz). Note that the theta and beta bands occur at lower frequencies under urethane anaesthesia compared to the awake state. Baseline values for LFP and fEPSP measures are shown in Table S1.

### Brain temperature recording

Brain temperature was measured using a Wheatstone bridge amplifier circuit to measure resistance changes in the thermistor located in a homotopic location to the recording electrode in the right hippocampus. Data were logged using the same NI-6003 acquisition device that was used to capture LFP activity, and calibrated by placing the thermistor in a water bath and recording the response across a range of temperatures. After capture, mean brain temperature values were averaged over successive 2-min recording periods throughout each experiment.

In the subset of saline-treated animals implanted with temperature probes, once the main experiment was completed, brain temperature was passively raised by removing the rectal temperature probe while LFP, fEPSP, and brain temperature recording were continued. This allowed an estimation of the expected change in fEPSP and LFP measures resulting from changes in brain temperature alone. For each animal, regression lines were calculated relating temperature increase to the measure of interest (Fig. S3 & S4). The mean slope of the relationship was used to calculate a predicted temperature-associated change in LFP and fEPSP measures for animals comprising the tianeptine- and buprenorphine-treatment groups in the main experiment. These predicted values were then compared with the observed values (see Table S2).

### Data analysis and statistics

Numerical data were analysed using Microsoft Excel and SPSS. Graphs were prepared using Excel, SigmaPlot, and Adobe Illustrator. Data are displayed as mean ± standard error of the mean (SEM), and individual data points are shown when bar graphs are presented. Where comparisons involved more than two groups, an ANOVA was performed followed, where appropriate, by post-hoc Tukey’s HSD pairwise comparisons. Comparisons between two groups comprised independent samples t-tests. One-sample t-tests with Bonferroni correction for multiple comparisons were used to compare the means of single groups with baseline values.

## Results

As reported previously, spontaneous hippocampal LFP activity under urethane anaesthesia was dominated by a slow-wave-sleep-like pattern comprising large amplitude irregular activity (LIA), with occasional episodes of theta activity reminiscent of REM sleep (e.g. Martin et al., 2019). Fig. 2A(i) shows a time-frequency plot of spectral power before and after the injection of vehicle (saline; n = 12); mean spectral power as a function of frequency, plotted before and 20-30 min after injection, is shown in (ii). Note the steep fall in power with increasing frequency and the absence of a lasting effect of injection. Figure 2A(iii) shows a time-frequency plot of the change in spectral power, calculated by subtracting the power at each time point and frequency bin from the corresponding mean value over the 10-min period before injection. The mean change in spectral power 20-30 min after injection is shown in (iv), and the inset shows a sample of LFP recording 20-30 min after injection, illustrating the characteristic LIA pattern of baseline activity.

**Fig. 2.**
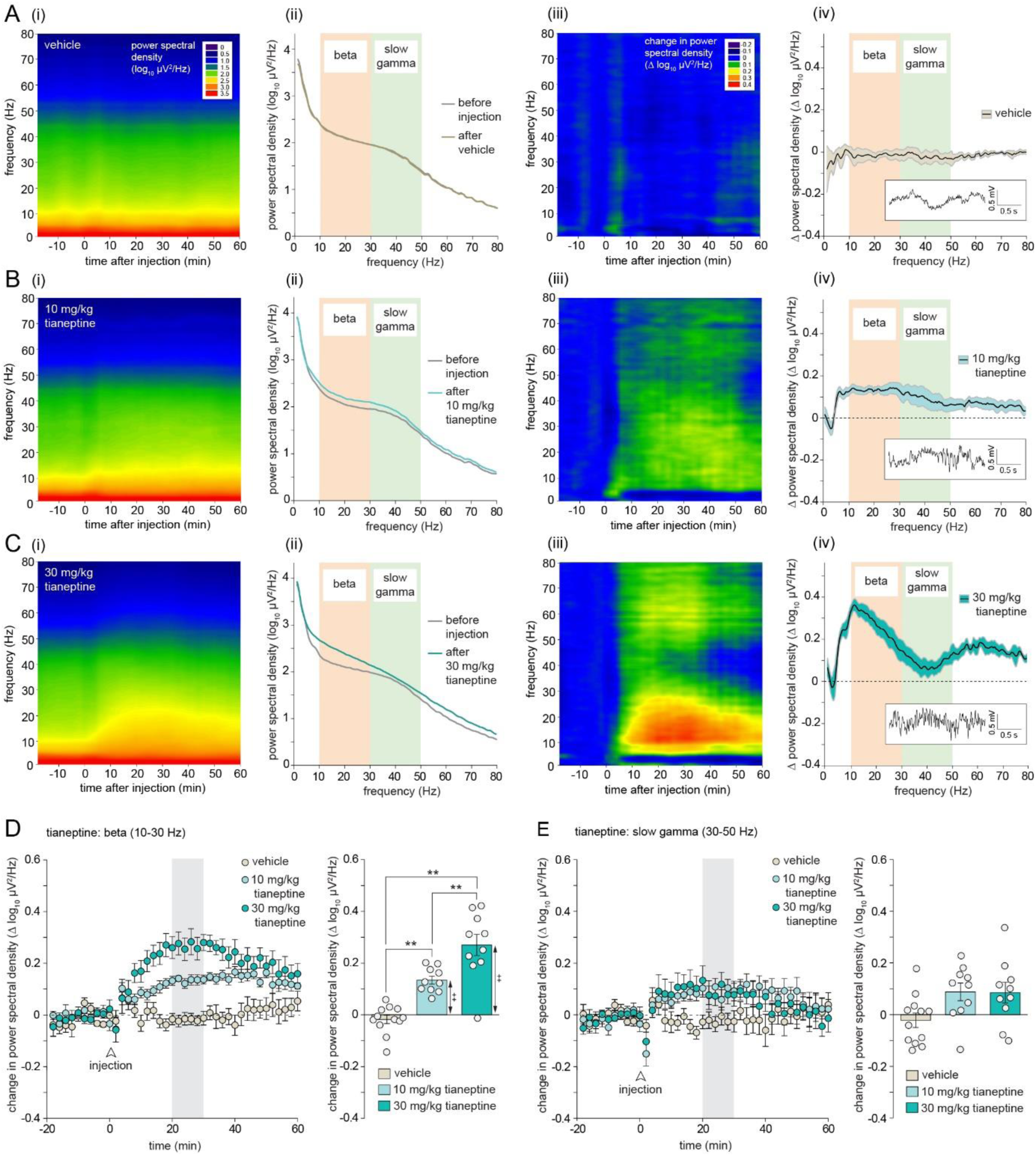
Tianeptine causes a dose-dependent increase in hippocampal beta-frequency oscillations. (A) (i) Time-frequency plot of mean spectral power after vehicle injection (n = 12). (ii) Mean spectral power over the 10-min baseline period before vehicle injection and 20-30 min after. (iii) Mean change in spectral power after vehicle injection. (iv) Mean change in spectral power 20-30 min after vehicle injection relative to baseline. The black lines indicate the means, and the filled areas bounded by grey lines indicate ± 1 SEM. The inset shows a sample of LFP activity recorded from a single rat ∼30 min after saline injection. (B) (i) Time-frequency plot of mean spectral power after 10 mg/kg tianeptine injection (n = 10). (ii) Mean spectral power over the 10-min baseline period before injection and 20-30 min after injection of 10 mg/kg tianeptine. (iii) Mean change in spectral power after 10 mg/kg tianeptine injection. (iv) Mean change in spectral power 20-30 min after 10 mg/kg tianeptine injection relative to baseline. The inset shows a sample of LFP activity recorded from a single rat ∼30 min after 10 mg/kg tianeptine injection. (C) (i) Time-frequency plot of mean spectral power after 30 mg/kg tianeptine injection (n = 10). (ii) Mean spectral power over the 10-min baseline period before 30 mg/kg tianeptine injection and 20-30 min after. (iii) Mean change in spectral power after 30 mg/kg tianeptine injection. (iv) Mean change in spectral power 20-30 min after 30 mg/kg tianeptine injection relative to baseline. The inset shows a sample of LFP activity recorded from a single rat ∼30 min after 30 mg/kg tianeptine injection. (D) Left-hand panel: time-course of tianeptine-induced changes in beta-frequency power normalised to the pre-injection baseline; right-hand panel: mean change in beta-frequency power 20-30 min after injection relative to baseline (**p < 0.01 post-hoc pairwise comparisons; ++p < 0.01 one-sample t-tests). (E) Tianeptine-induced change in slow-gamma-frequency power; other details as for D.

Fig. 2B shows the effects of 10 mg/kg tianeptine (n = 10) plotted in the same way. Analysis of the change in spectral power (panels iii and iv) revealed a broad increase in spectral power that was maximal in the beta-frequency range, illustrated by the appearance of rhythmic beta-frequency oscillations in the LFP trace (see inset, panel iv). Fig. 2C shows the spectral changes caused by 30 mg/kg tianeptine (n = 10). This dose produced a more pronounced increase in the low-beta-frequency range, and a second, less-pronounced increased in fast gamma activity. Analysis of the time-course of the tianeptine-induced increase in beta activity revealed a gradual increase in power that peaked 20-30 min after injection at the 30 mg/kg dose (Fig. 2D; left-hand panel). Analysis of the mean change in power 20-30 min after injection (Fig. 2D; right-hand panel) indicated a dose-dependent increase in beta power; an ANOVA revealed a significant main effect of tianeptine dose [F(2,29) = 32.2; p < 0.001], and post-hoc Tukey’s HSD pairwise comparisons revealed significant differences between the effects of all doses (vehicle verses 10 mg/kg & 30 mg/kg tianeptine: p < 0.001 in both cases; 10 mg/kg tianeptine versus 30 mg/kg tianeptine: p = 0.0003). Significant increases relative to baseline were observed at both 10 mg/kg (p < 0.0003) and 30 mg/kg doses (p = 0.0003; one-sample t-tests with Bonferroni correction for multiple comparisons). The effects of tianeptine on slow gamma power were modest (Fig. 2E); an ANOVA of mean power 20-30 min after injection (Fig. 2E; right hand panel) revealed a significant effect of increasing tianeptine dose [F(2,29) = 3.73; p = 0.036], but post-hoc Tukey’s HSD pairwise comparisons did not reveal any significant pairwise differences, and none of the groups differed significantly from baseline. An analysis of changes in theta and fast gamma activity is presented in Fig. S2A&B.

The increase in beta power induced by 30 mg/kg tianeptine was completely blocked by co-administration of 1 mg/kg naloxone (Fig. 3A). Injection of naloxone alone had no effect (Fig. 3B). The samples of LFP activity shown in the insets of panels (iv) indicate a characteristic LIA pattern in both cases. The time-course of changes in beta power in the tianeptine + naloxone, and naloxone only groups is shown in Fig. 3C (left-hand panel), with the 30 mg/kg tianeptine data from Fig. 1D included for comparison. An ANOVA of the mean beta power in the 3 groups 20-30 min after injection (Fig. 3C, right-hand panel) revealed a significant main effect of group [F(2,22) = 18.4; p < 0.001], with significantly lower power in both the tianeptine + naloxone and naloxone only groups relative to the 30 mg/kg tianeptine group (Tukey’s HSD; p < 0.001 in both cases); no significant change in beta power relative to baseline was observed in the tianeptine + naloxone, and naloxone only groups. An ANOVA of the mean slow gamma power in the 3 groups 20-30 min after injection (Fig. 3D, right-hand panel) did not reveal a significant group difference [F(2,22) = 2.50; p = 0.11]; no significant change in slow gamma power relative to baseline was observed in the tianeptine + naloxone, and naloxone only groups. Corresponding changes in theta and fast gamma activity are presented in Fig. S2C&D.

**Fig. 3.**
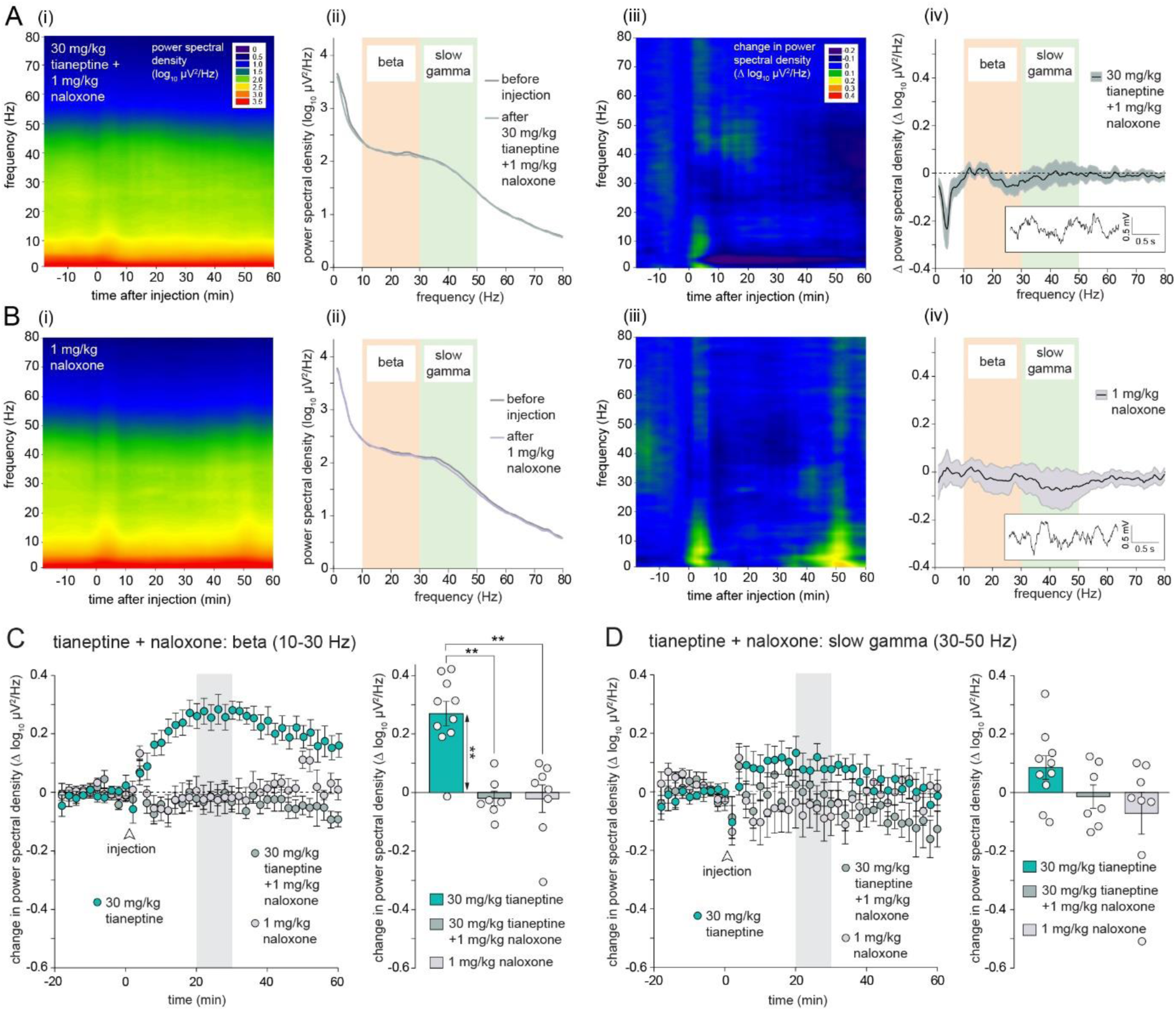
The tianeptine-induced increase in beta-frequency oscillations is blocked by naloxone. (A) (i) Time-frequency plot of mean spectral power after co-injection of 30 mg/kg tianeptine and 1 mg/kg naloxone (n = 7). (ii) Mean spectral power over the 10-min baseline period before tianeptine and naloxone injection and 20-30 min after. (iii) Mean change in spectral power after tianeptine and naloxone injection. (iv) Mean change in spectral power 20-30 min after tianeptine and naloxone injection relative to baseline. The inset shows a sample of LFP activity recorded from a single rat ∼30 min after tianeptine and naloxone injection. (B) (i) Time-frequency plot of mean spectral power after injection of 1 mg/kg naloxone alone (n = 8). (ii) Mean spectral power over the 10-min baseline period before naloxone injection and 20-30 min after. (iii) Mean change in spectral power after naloxone injection. (iv) Mean change in spectral power 20-30 min after naloxone injection relative to baseline. The inset shows a sample of LFP activity recorded from a single rat ∼30 min after naloxone injection. (C) Left-hand panel: time-course of changes in beta-frequency power after co-administration of tianeptine and naloxone versus naloxone alone. The response to 30 mg/kg tianeptine only is plotted for comparison (reproduced from fig. 2D); right-hand panel: mean change in beta-frequency power in the same 3 groups 20-30 min after injection relative to baseline (**p < 0.01 post-hoc pairwise comparisons; ++p < 0.01 one-sample t-tests). (D) Left-hand panel: time-course of changes in slow gamma power after administration of tianeptine + naloxone, naloxone only, and tianeptine only; right-hand panel: mean change in beta-frequency power in the same 3 groups 20-30 min after injection relative to baseline.

Fig. 4 shows the corresponding changes in fEPSPs in the same animals for which LFP data is shown in Fig. 2 & 3. Examples of fEPSPs from individual rats after injection of vehicle or 30 mg/kg tianeptine are shown in Fig. 4A. Note the slight increase in the slope and peak amplitude after tianeptine injection, and the reduction in amplitude of the positive overshoot portion of the fEPSP tail. An ANOVA of the mean slope increase 20-30 min after injection in the vehicle and tianeptine groups (Fig. 4B) revealed a main effect of drug dose [F(2,29) = 3.83; p = 0.034]; post-hoc pairwise comparisons (Tukey’s HSD) revealed a significant difference between saline and 30 mg/kg tianeptine groups (p = 0.027), and a significant increase relative to baseline was evident at the 30 mg/kg dose (p = 0.003; one-sample t-test with Bonferroni correction). The amplitude of the tail of the fEPSP 20-30 min after injection (Fig. 4C; see Fig. S4) showed a dose dependent decrease, indicated by the main effect of drug dose [F(2,29) = 11.9; p < 0.001]; post-hoc pairwise comparisons (Tukey’s HSD) revealed a significant difference between vehicle and 30 mg/kg tianeptine groups (p < 0.001) and between 10 mg/kg and 30 mg/kg tianeptine groups (p = 0.015). Both 10 mg/kg and 30 mg/kg tianeptine groups showed significant decreases from baseline (p = 0.021 & p = 0.0007 respectively; one-sample t-tests with Bonferroni correction).

**Fig. 4.**
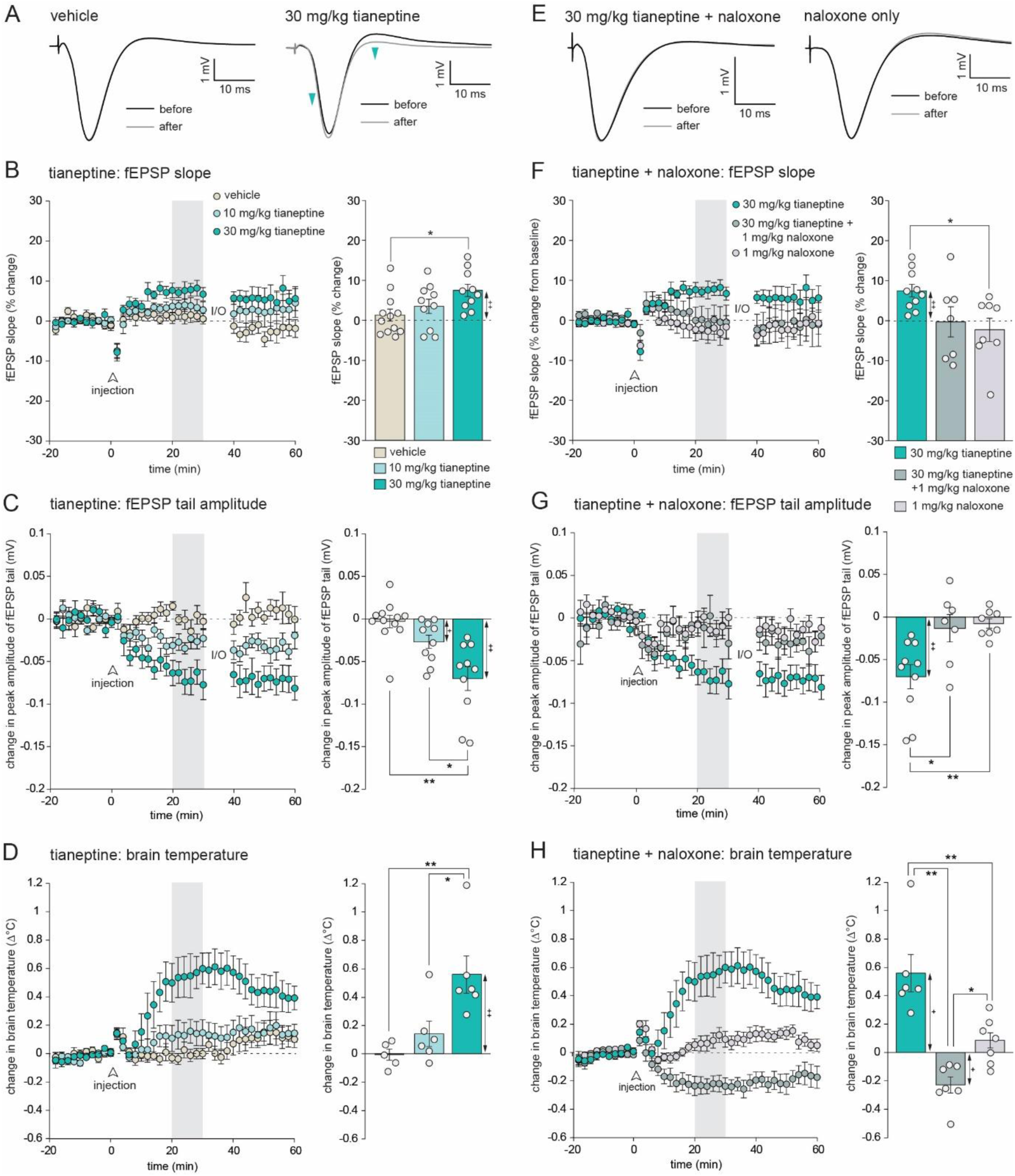
Tianeptine causes a dose-dependent increase in fEPSP slope associated with an increase in brain temperature. (A) Examples of fEPSPs recorded before and ∼30 min after the infusion of vehicle and 30 mg/kg tianeptine. Blue arrowheads indicate the modest increase in fEPSP slope and amplitude, and the reduction in amplitude of the ‘tail’ of the fEPSP after 30 mg/kg tianeptine. (B) Left-hand panel: time course of the mean increase in fEPSP slope after injection of saline (n = 12) or tianeptine at 10 mg/kg (n = 10) and 30 mg/kg (n = 10). The missing portion of data corresponds to an input-ouput (I/O) curve that was recorded starting 30 min after tianeptine injection. Right-hand panel: mean fEPSP slope 20-30 min after injection normalised to baseline. (C) Left-hand panel: time course of the mean decrease in fEPSP tail amplitude after injection of saline or tianeptine at 10 and 30 mg/kg. Right-hand panel: mean fEPSP tail amplitude 20-30 min after injection. (D) Time course of the mean increase in brain temperature after injection of saline or tianeptine at 10 and 30 mg/kg. Right-hand panel: mean change in brain temperature 20-30 min after injection. (E) Examples of fEPSPs recorded before and ∼30 min after the administration of tianeptine + naloxone, or naloxone only. (F) Left-hand panel: time course of the mean increase in fEPSP slope after injection of tianeptine + naloxone (n = 7) or naloxone only (n = 8); the tianeptine group from B is reproduced for comparison. Right-hand panel: mean fEPSP slope 20-30 min after injection normalised to baseline. (G) Left-hand panel: time course of the mean decrease in fEPSP tail amplitude after injection of tianeptine + naloxone, naloxone only, and tianeptine only. Right-hand panel: mean change in fEPSP tail amplitude 20-30 min after injection. (H) Left-hand panel: time course of the change in brain temperature after injection of tianeptine + naloxone, naloxone, and tianeptine. Right-hand panel: mean change in brain temperature 20-30 min after injection. **p < 0.01, *p < 0.05: post-hoc pairwise comparisons; ++p < 0.01, +p < 0.05: one-sample t-tests.

Fig. 4D shows the effect of tianeptine on brain temperature in the subsets of animals implanted with temperature probes (vehicle: n = 5; 10 mg/kg tianeptine: n = 6; 30 mg/kg tianeptine: n = 6). At the 30 mg/kg dose, tianeptine caused a pronounced temperature increase averaging ∼0.6°C. An ANOVA of the mean increase 20-30 min after injection revealed a significant main effect of tianeptine dose [F(2,14) = 8.56; p = 0.004]; post-hoc pairwise comparisons (Tukey’s HSD) revealed a significant difference between vehicle and 30 mg/kg tianeptine groups (p = 0.004) and between 10 mg/kg and 30 mg/kg tianeptine groups (p = 0.022). The temperature rise in the 30 mg/kg tianeptine group was significantly increased relative to baseline (p < 0.024; one-sample t-test with Bonferroni correction).

Co-administration of 30 mg/kg tianeptine with 1 mg/kg naloxone (n = 7) completely blocked the fEPSP slope increase (Fig. 4 E & F). Examples of fEPSPs before and after the injection of 30 mg/kg tianeptine + 1 mg/kg naloxone and naloxone only (Fig. 4E; n = 8) illustrate the absence of any drug-related changes. An ANOVA of the mean increase 20-30 min after injection in the 30 mg/kg tianeptine, 30 mg/kg tianeptine + 1 mg/kg naloxone, and naloxone only groups (Fig. 4F) revealed a main effect of group [F(2,22) = 4.16; p = 0.029], and a significant difference between the 30 mg/kg tianeptine and 30 mg/kg tianeptine + 1 mg/kg naloxone groups (p < 0.034; post-hoc pairwise comparison; Tukey’s HSD). No significant changes from baseline were observed in the 30 mg/kg tianeptine + 1 mg/kg naloxone, and naloxone only groups. Naloxone also completely blocked the tianeptine-induced decrease in the fEPSP tail (Fig. 4G). An ANOVA of the mean increase 20-30 min after injection in the 30 mg/kg tianeptine, 30 mg/kg tianeptine + 1 mg/kg naloxone, and naloxone only groups revealed a main effect of group [F(2,22) = 7.78; p = 0.003]; post-hoc pairwise comparisons (Tukey’s HSD) indicated a significant difference between the 30 mg/kg tianeptine and the two other groups: 30 mg/kg tianeptine + 1 mg/kg naloxone (p < 0.014) and naloxone only (p = 0.005). No significant changes from baseline in the tail amplitude measure were observed in the 30 mg/kg tianeptine + 1 mg/kg naloxone, and naloxone only groups.

Naloxone alone (n = 8) had no effect on brain temperature, but co-administration of 30 mg/kg tianeptine + 1mg/kg naloxone (n = 7) resulted in a slight drop in temperature (Fig. 4H). The reasons for this are unclear, but may reflect the unmasking of an additional opioid-receptor-independent temperature-lowering effect of tianeptine. An ANOVA of the temperature change 20-30 min after injection in the 30 mg/kg tianeptine, 30 mg/kg tianeptine + naloxone, and naloxone only groups revealed a main effect of group [F(2,18 = 22.5; p < 0.001]; post-hoc pairwise comparisons (Tukey’s HSD) revealed significant differences between all groups (30 mg/kg tianeptine versus 30 mg/kg tianeptine + 1 mg/kg naloxone: p < 0.001; 30 mg/kg tianeptine versus naloxone only: p = 0.002; 30 mg/kg tianeptine + 1 mg/kg naloxone versus naloxone only: p < 0.026). There was a significant increase in temperature relative to baseline in the 30 mg/kg tianeptine group (p = 0.024), and a significant decrease in the 30 mg/kg tianeptine + 1 mg/kg naloxone group (p = 0.018; one-sample t-test with Bonferroni correction).

Figure 5 shows the effects of another opioid receptor agonist with antidepressant actions, buprenorphine, on hippocampal LFP activity. Injection of 0.03 mg/kg buprenorphine caused a similar pattern of activity to tianeptine, with a peak in the low-beta-frequency range, and an additional smaller peak in the fast gamma range (Fig. 5A). These changes were completely blocked by the co-administration of 1 mg/kg naloxone (Fig. 5B), confirming their dependence on opioid receptor activation. Buprenorphine showed a slightly longer time-course of action compared to tianeptine, with the peak increase in beta power occurring ∼50 min after injection (Fig. 5C; left-hand panel), but for consistency with previous analyses, the mean power 20-30 min after injection was calculated; injection of buprenorphine produced an above-chance increase in beta power (p = 0.016; one-sample t-test with Bonferroni correction) that was significantly reduced when co-administered with naloxone [t(10) = 3.15; p = 0.01; independent samples t-test; Fig. 5C; right-hand panel]. There was also significantly higher slow gamma activity 20-30 min after injection in the buprenorphine group relative to the buprenorphine + naloxone group [t(10) = 2.44; p = 0.035; Fig. 5D]. An analysis of buprenorphine-induced changes in theta and fast gamma changes in presented in Supplementary Fig. S2E & F.

**Fig 5.**
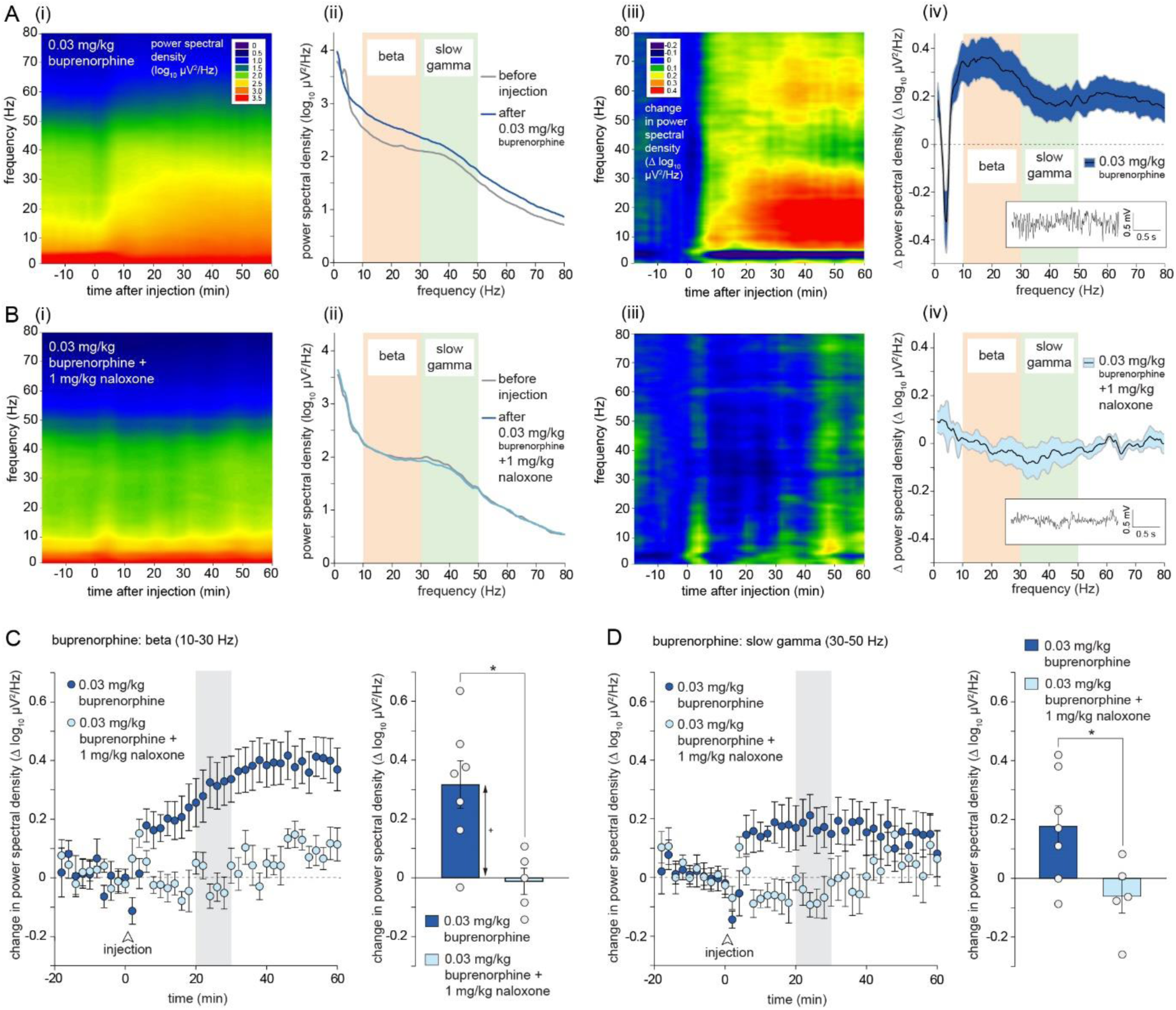
Buprenorphine causes a similar naloxone-sensitive increase in beta-frequency oscillations to tianeptine. (A) (i) Time-frequency plot of mean spectral power after injection of 0.03 mg/kg buprenorphine (n = 7). (ii) Mean spectral power over the 10-min baseline before buprenorphine administration and 20-30 min after. (iii) Mean change in spectral power after buprenorphine injection. (iv) Mean change in spectral power 20-30 min after buprenorphine injection relative to baseline. The inset shows a segment of LFP activity recorded from a single rat ∼30 min injection. Note the presence of rhythmic beta-frequency oscillations. (B) Time-frequency plot of mean spectral power after co-injection of 0.03 mg/kg buprenorphine and 1 mg/kg naloxone (n = 5). (ii) Mean spectral power over the 10-min baseline period before buprenorphine and naloxone injection and 20-30 min after. (iii) Time-frequency plot of the mean change in spectral power after buprenorphine and naloxone injection. (iv) Mean change in spectral power 20-30 min after buprenorphine and naloxone injection relative to baseline. The inset shows a sample of LFP activity recorded ∼30 min after injection. (C) Left-hand panel: time-course of changes in beta-frequency power after administration of buprenorphine versus buprenorphine + naloxone; right-hand panel: mean change in beta-frequency power 20-30 min after injection relative to baseline. (D) Left-hand panel: time-course of changes in slow gamma power after administration of buprenorphine versus buprenorphine + naloxone; right-hand panel: mean change in slow gamma power 20-30 min after injection relative to baseline. *p < 0.05: independent samples t-test; +p < 0.05; one-sample t-test.

The fact that buprenorphine and tianeptine caused increases in brain temperature raises the possibility that the fEPSP and LFP changes caused by these drugs are a direct consequence of this temperature change. To assess this, we examined the relationship between brain temperature and electrophysiological measures in a subset of vehicle-treated animals once the main experiment was finished (Fig. S3 & S4). Using this data, we calculated predicted changes in fEPSP and LFP measures based on the observed change in brain temperature in the subsets of tianeptine- and buprenorphine-injected animals implanted with temperature probes. This analysis revealed that the observed and expected changes in fEPSP slope did not differ in any group (Table S2A), i.e. the slight tianeptine- and buprenorphine-induced increase in fEPSP slope was fully predicted by the increase in brain temperature. However, the tianeptine-induced reduction in amplitude of the fEPSP tail was significantly larger than predicted based on the temperature rise alone (Table S2B).

Figure 6 shows the effects of ketamine, a fast-acting antidepressant with a different mechanism of action. Consistent with previous reports, 10 mg/kg ketamine caused an increase in slow gamma power, but a decrease in frequencies below ∼20 Hz (Fig. 6A), i.e. a profile of LFP changes distinct from that produced by tianeptine and buprenorphine. Ketamine-induced gamma oscillations were unaffected by co-administration of naloxone (Fig. 6B). Ketamine had no overall effect on mean power in the beta-frequency band 20-30 min after injection (t(17) = 1.65; p = 0.118; Fig. 6C) since power in the low beta range was reduced while power at the upper end of the range was increased. The peak increase in slow gamma power occurred ∼20-30 min after injection, and mean power in this range was significantly increased in both the ketamine and ketamine + naloxone groups (p = 0.0012 and 0.0034 respectively; one-sample t-tests with Bonferroni correction), with no difference between the two groups [t(17) = 2.08; p = 0.053]. An analysis of ketamine-induced changes in theta and fast gamma changes is presented in Supplementary Fig. S2G & H.

**Fig. 6.**
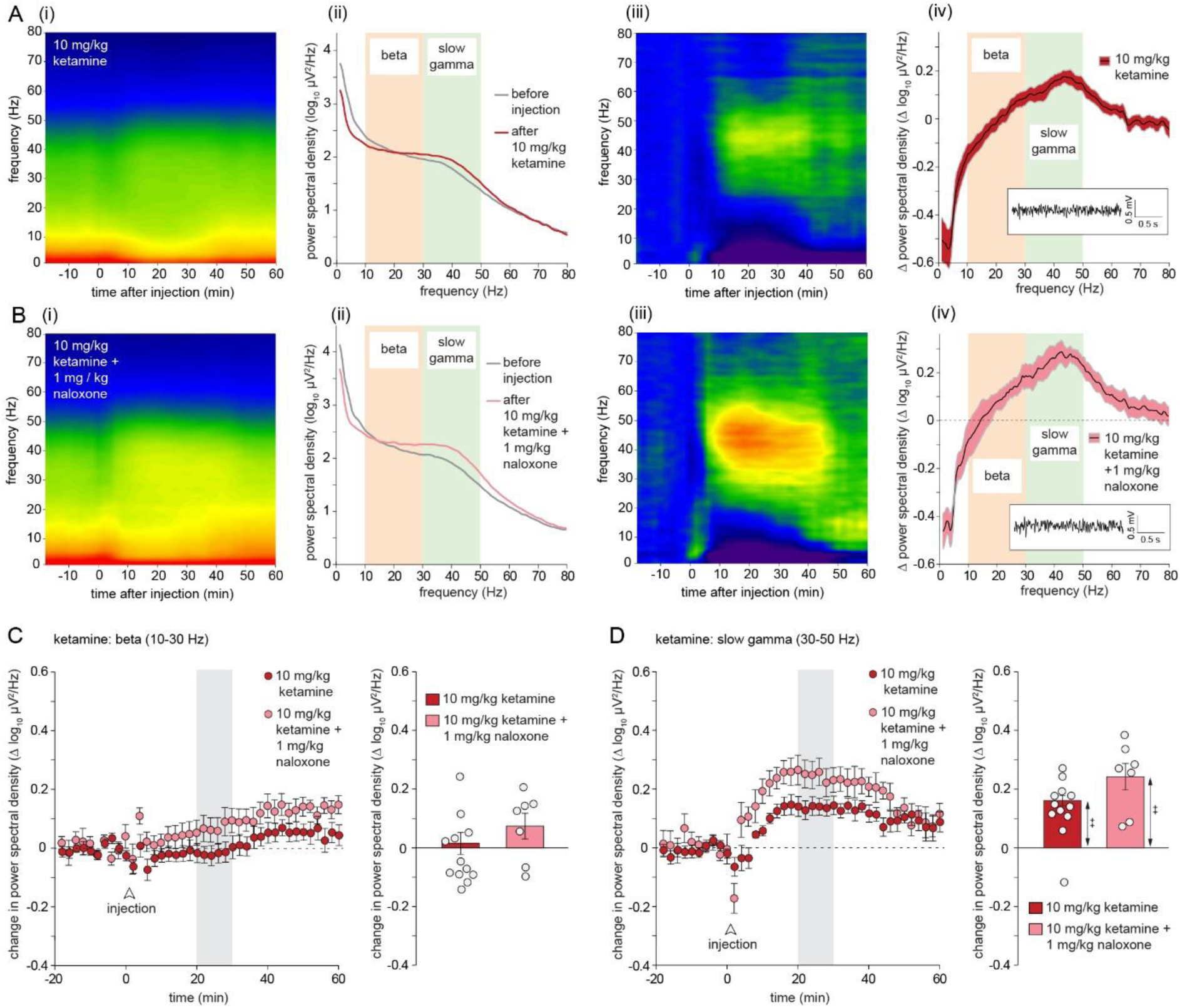
Ketamine causes a distinct pattern of slow gamma oscillations that is insensitive to naloxone. (A) (i) Time-frequency plot of mean spectral power after injection of 10 mg/kg ketamine (n = 12). (ii) Mean spectral power over the 10-min baseline before ketamine administration and 20-30 min after. (iii) Time-frequency plot of mean change in spectral power after ketamine injection. (iv) Mean change in spectral power 20-30 min after ketamine injection relative to baseline. The inset shows a segment of LFP activity recorded from a single rat ∼30 min injection, showing the presence of gamma-frequency oscillations. (B) Time-frequency plot of mean spectral power after co-injection of ketamine and 1 mg/kg naloxone (n = 7). (ii) Time-frequency plot of mean spectral power over the 10-min baseline period before ketamine + naloxone injection and 20-30 min after. (iii) Time-frequency plot of the mean change in spectral power after ketamine + naloxone injection. (iv) Mean change in spectral power 20-30 min after ketamine + naloxone injection relative to baseline. The inset shows a sample of LFP activity recorded ∼30 min after injection, showing the presence of gamma-frequency oscillations. (C) Left-hand panel: time-course of changes in beta-frequency power after administration of ketamine and ketamine + naloxone; right-hand panel: mean change in beta-frequency power 20-30 min after injection relative to baseline. (D) Left-hand panel: time-course of changes in slow gamma power after administration of ketamine and ketamine + naloxone; right-hand panel: mean change in slow gamma power 20-30 min after injection relative to baseline (++p< 0.01; one-sample t-tests).

Fig. 7 shows fEPSP changes following buprenorphine and ketamine injection, corresponding to the LFP data shown in Fig. 5 & 6. The effect of buprenorphine on fEPSPs was qualitatively similar to that of tianeptine. Examples of fEPSPs from a single rat after injection of 0.03 mg/kg buprenorphine are shown in Fig. 7A. Note the slight increase in the slope and peak amplitude after buprenorphine injection, and the slight reduction in amplitude of the positive overshoot portion of the fEPSP tail (left-hand waveforms; blue arrowheads). In this example, the changes were reduced by co-administration of naloxone, illustrated by the right-hand waveforms. Overall, buprenorphine caused a modest numerical increase in fEPSP slope, but mean slope 20-30 min after injection did not differ in the buprenorphine and buprenorphine + naloxone groups [t(10) = 1.66; p = 0.13], and one-sample t-tests with Bonferroni corrections did not indicate significant increases relative to baseline in either group. Likewise, buprenorphine caused a small decrease in the amplitude of the fEPSP tail that was numerically smaller after naloxone administration, but the difference between the groups did not reach significance 20-30 min after injection [t(10) = 1.76; p = 0.10; Fig. 5 C], and one-sample t-tests with Bonferroni corrections did not indicate significant increases relative to baseline in either group. However, brain temperature was significantly increased 20-30 min after buprenorphine injection (p = 0.0004; one-sample t-test with Bonferroni correction); this increase was significantly reduced in the presence of naloxone [t(10) = 4.15; p = 0.002; Fig. 7D]. As with tianeptine, the small increase in fEPSP after buprenorphine injection was fully predicted by the rise in brain temperature, but the temperature increase cannot account for the change in beta power (see Table S2 for full analysis).

**Fig. 7.**
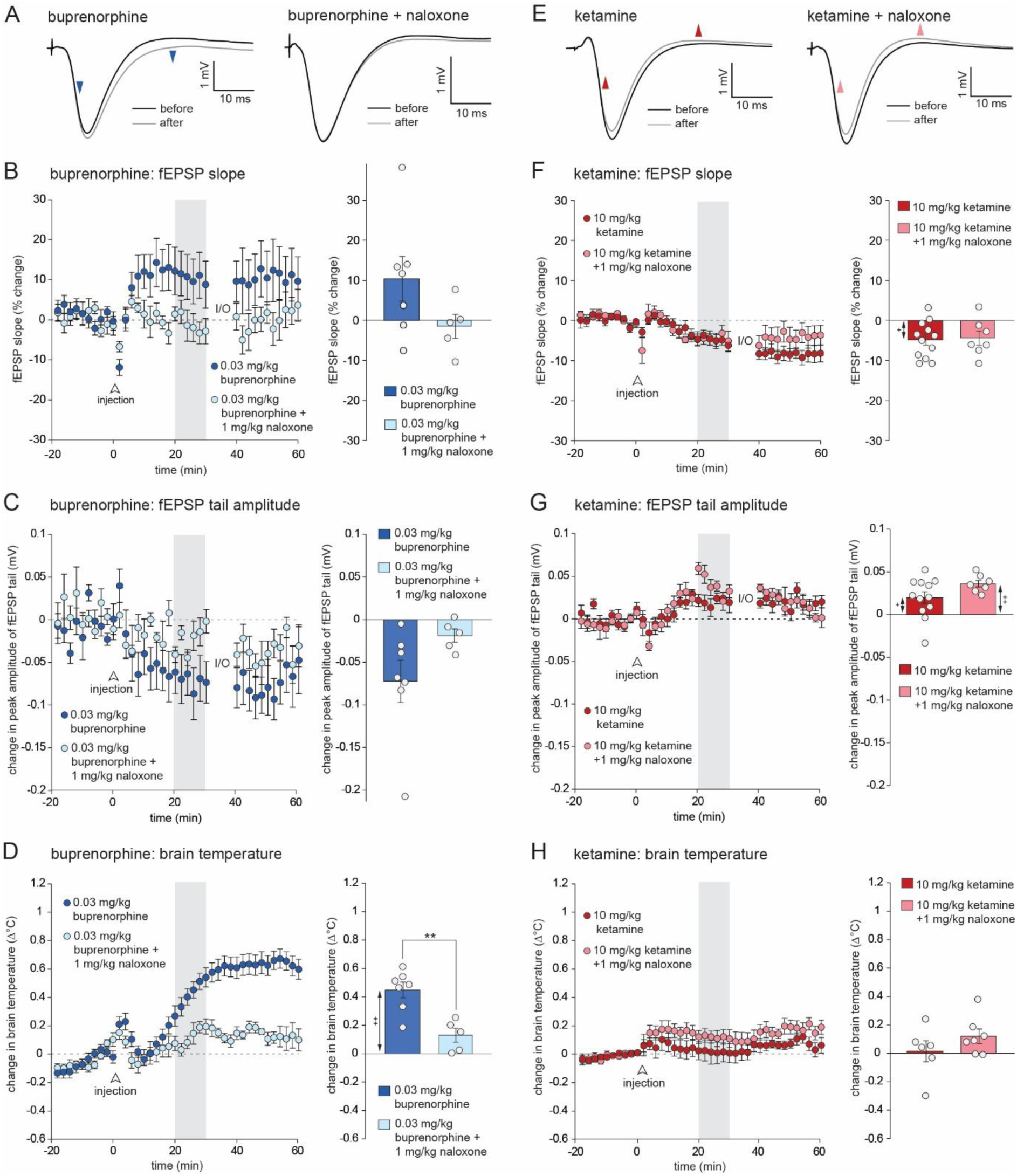
Buprenorphine, but not ketamine, causes similar fEPSP and brain temperature changes to tianeptine. (A) Examples of fEPSPs recorded before and ∼30 min after the infusion of 0.03 mg/kg buprenorphine and 0.03 mg/kg buprenorphine + 1 mg/kg naloxone. Blue arrowheads indicate the modest increase in fEPSP slope and amplitude, and the reduction in amplitude of the ‘tail’ of the fEPSP after buprenorphine. (B) Left-hand panel: time course of the mean increase in fEPSP slope after injection of 0.03 mg/kg buprenorphine (n = 7) or buprenorphine + naloxone (n = 5); right-hand panel: mean fEPSP slope 20-30 min after injection normalised to baseline. (C) Left-hand panel: time course of the mean decrease in fEPSP tail amplitude after injection of buprenorphine versus buprenorphine + naloxone. Right-hand panel: mean fEPSP tail amplitude 20-30 min after injection. (D) Time course of the mean increase in brain temperature after injection of buprenorphine versus buprenorphine + naloxone; right-hand panel: mean change in brain temperature 20-30 min after injection. (E) Examples of fEPSPs recorded before and ∼30 min after the administration of ketamine and ketamine + naloxone; note the small reduction in the slope and peak amplitude of the fEPSP in each case. (F) Left-hand panel: time course of the mean increase in fEPSP slope after injection of 10 mg/kg ketamine (n = 12) and 10 mg/kg ketamine + 1 mg/kg naloxone (n = 7); right-hand panel: mean fEPSP slope 20-30 min after injection normalised to baseline. (G) Left-hand panel: time course of the mean increase in fEPSP tail amplitude after injection of ketamine versus ketamine + naloxone; right-hand panel: mean change in fEPSP tail amplitude 20-30 min after injection. (H) Left-hand panel: time course of the change in brain temperature after injection of ketamine versus ketamine + naloxone; right-hand panel: mean change in brain temperature 20-30 min after injection. **p < 0.01: independent samples t-test; ++p < 0.01, +p < 0.05: one-sample t-tests.

Examples of fEPSPs from rats injected with 10 mg/kg ketamine, or 10 mg/kg ketamine + 1 mg/kg naloxone are shown in Fig. 7E. In both cases, a small reduction in the amplitude and slope of the fEPSP is seen, together with a small increase in the positive amplitude of the tail. The mean time course of these effects is shown in Fig. 7F; there was no difference in fEPSP slope between ketamine and ketamine + naloxone groups 20-30 min after injection [t(17) = 0.21; p = 0.84], but there was a significant reduction relative to baseline in the ketamine group (p = 0.013; one-sample t-test with Bonferroni correction). The two groups also did not differ in the amplitude of the fEPSP tail after injection [t(17) = 1.73; p = 0.10; Fig. 7G], but both groups showed a significant increase in the amplitude of this measure (ketamine: p = 0.027; ketamine + naloxone: p < 0.0002; one-sample t-tests with Bonferroni correction). No significant increase in brain temperature was observed 20-30 min after injection of ketamine or ketamine + naloxone (one-sample t-tests with Bonferroni correction; Fig. 7H), and the two groups did not differ from each other [t(11) = 1.22; p = 0.25].

## Discussion

Our key finding is that tianeptine causes a dose-dependent increase in hippocampal LFP oscillations at the lower end of the beta-frequency range. This effect was completely abolished by co-administration of the opioid receptor antagonist naloxone. Another opioid receptor agonist with antidepressant actions, buprenorphine, caused a similar naloxone-sensitive increase in beta-frequency activity. Naloxone alone had no effect on LFP power, suggesting that endogenous opioid-receptor activity does not contribute to oscillatory activity under these conditions. Ketamine, in contrast, produced a distinct profile of LFP changes, characterised by a decrease in power below 20 Hz and an increase in slow gamma power that was insensitive to naloxone. Previous EEG studies have reported tianeptine-induced increases in cortical beta-frequency activity in humans (Saletu et al., 1996) and prefrontal gamma in mice (Rozov et al., 2021). Based on these findings and those of the present study, tianeptine belongs to a growing list of antidepressants that enhance beta / gamma oscillations in the hippocampus or neocortex, a phenomenon that has been proposed as a marker for antidepressant efficacy (Fitzgerald & Watson, 2019; Gilbert & Zarate Jr 2020). However, not all compounds that increase power in the beta and gamma ranges have antidepressant actions; for example, benzodiazepines enhance beta and slow-gamma oscillations (Lambert et al., 2023), but are not effective as antidepressants despite their anxiolytic actions (Lim et al., 2020). Nonetheless, while not sufficient for antidepressant efficacy, it is possible that this pattern of activity plays a necessary role in the actions of some antidepressant drugs.

Tianeptine also caused a modest increase of CA1 evoked fEPSPs. At face value, this effect would seem consistent with in vitro evidence that tianeptine can enhance AMPA-receptor-mediated fast synaptic transmission (Kole et al., 2002; Szegedi et al., 2011; Pillai et al., 2012; Zhang et al., 2013 & 2018). However, the fEPSP increase was associated with an increase in brain temperature of ∼0.6°C at a dose of 30 mg/kg. Analysis of the relationship between fEPSP slope and temperature in a subset of animals revealed that the observed increase in fEPSPs was fully predicted by the observed temperature increase. In contrast, tianeptine-induced LFP changes could not be explained by the temperature increase alone; in fact, analysis of the relationship between brain temperature and beta power predicts a modest decrease, rather than the increase observed. Both the temperature and fEPSP changes caused by tianeptine were blocked by naloxone, consistent with previous reports of brain and core temperature increases following the administration of morphine (Solis et al., 2018).

Tianeptine’s peak effect on beta-frequency activity occurred 20-30 min after injection of a 30 mg/kg dose. This would appear to parallel the pharmacokinetic profile of tianeptine’s principal metabolite MC5 more closely than that of tianeptine itself. Brain concentrations of tianeptine peak within 5 min of systemic injection in rats and mice and fall rapidly thereafter; in contrast, MC5 concentrations peak after ∼30 min and exhibit a much longer half-life (Couet et al., 1990; Samuels et al., 2017). Both tianeptine and MC5 are opioid receptor agonists with similar efficacy and potency at the µ-OR, although both have lower potency at the δ-OR (Gassaway et al., 2014; Samuels et al., 2017). However, confirmation of the relative roles of tianeptine versus MC5 in the oscillatory activity reported here will require a direct comparison of the effects of the two compounds. It is also possible that the time-course of LFP changes may reflect dynamic responses to the actions of a drug that are not solely determined by its concentration at a specific time.

The identity and location of the opioid receptors responsible for tianeptine or MC5’s enhancement of beta oscillations cannot be determined definitively based on the present data alone. Previous reports indicate that a canonical opioid, morphine, also a non-subtype-selective agonist, can enhance hippocampal and striatal gamma oscillations (Sakae & Martin, 2019; Reakkamnuan et al., 2023). However, our preliminary data indicate that activation of the µ-opioid receptor is central to tianeptine’s LFP effects (Trigo et al., 2022). A possibility is that the activation of µ-opioid receptors on hippocampal interneurons causes a reduction in GABA release, or interneuron firing, resulting in a disinhibition of pyramidal neurons. This is a well characterised action of µ-opioid receptor agonists, and reflects a G-protein-dependent inhibition of voltage-activated calcium channels leading to decreased GABA release, and / or the opening of G-protein-coupled inwardly rectifying potassium (GIRK) channels leading to a reduction in interneuron excitability and firing rate (see Nam et al., 2021). The possibility of disinhibition is consistent with our observation that tianeptine causes a decrease in the amplitude of the overshoot portion of the fEPSP tail. Recruitment of Schaffer collateral afferents results in a dual component fEPSP, comprising an excitatory component mediated by activation of glutamatergic synapses and a slower opposing component resulting from the feedforward recruitment of inhibitory interneurons (Alger & Nicoll, 1982; Arai et al., 1995; Karnup & Stelzer, 1999). A reduction in the inhibitory component, for example caused by a µ-opioid-receptor-dependent reduction in interneuron firing and / or GABA release, might account for the observed reduction in the amplitude of the tail portion of the fEPSP (see Supplementary Fig. S4). A disinhibition of principal neurons caused by the inhibition of NMDA receptors on GABAergic interneurons has been proposed as a mechanism for ketamine-induced gamma oscillations (e.g. Neymotin et al., 2011), although a recent modelling study indicates that the selective inhibition of NMDA receptors on inhibitory interneurons is unnecessary for the generation of gamma (Adam et al., 2024). Disinhibition of hippocampal pyramidal cells by pharmacological inhibition of GABA_A_ receptors also causes changes in LFP activity—for example, local infusion of picrotoxin into the ventral hippocampus of isoflurane-anaesthetised rats causes an increase in spectral power at frequencies below 20 Hz (Gwilt et al., 2020). This is reminiscent of the low-beta frequency changes caused by tianeptine in the present study, but a direct comparison is challenging owing to the distinct background LFP states produced by urethane versus isoflurane anaesthesia (Gwilt et al., 2020). We are currently conducting a more detailed characterisation of tianeptine’s effects on inhibitory neurotransmission and its consequences for hippocampal network activity.

An additional question concerns the regional site of action of tianeptine in the current study. Since tianeptine was administered systemically, the whole brain will have been exposed to the drug. It is likely that the observed oscillations originate in the hippocampus itself—slow gamma oscillations (∼25-50 Hz), for example, can originate in CA3, leading to coupled oscillations in CA1 (Colgin et al., 2015). However, it is possible that associated brain regions such as the medial septum or entorhinal cortex may play a major or contributory role in the observed LFP changes. Another potential site of action for tianeptine is the ventral tegmental area (VTA). Activation of µ-opioid receptors located on inhibitory interneurons of the VTA causes a disinhibition of dopaminergic neurons (Bull et al., 2017); these neurons project to the ventral striatum and other brain regions including the hippocampus; however, the latter pathway primarily targets the ventral hippocampus (Takeuchi et al., 2016; Tse, Privitera et al., 2023), whereas in the present study electrodes were placed in the dorsal part of the structure.

Although the systems-level mechanisms of tianeptine’s antidepressant actions are not fully understood, including the role of changes in network activity as reported here, there is considerable evidence regarding the cellular and neuronal mechanisms involved in tianeptine’s antidepressant effects. Regarding the role of opioid-receptors, a detailed series of experiments has revealed that µ-opioid receptor activation on somatostatin-positive interneurons of the ventral hippocampus is necessary for the antidepressant actions of tianeptine in mice (Han et al., 2022). Earlier work has also implicated an enhancement of hippocampal AMPA receptor function in the antidepressant actions of tianeptine. Tianeptine increases the phosphorylation of the GluA1 subunit at Ser831 and Ser 845 sites, and the introduction of a point mutation that prevents phosphorylation at both of these sites abolished tianeptine’s antidepressant-like effects in the tail suspension task (Svenningson et al., 2007). Taking these studies together, it is possible that both a reduction in presynaptic inhibitory transmission, and an enhancement of postsynaptic AMPA-receptor-mediated excitation are both required for tianeptine’s antidepressant actions, shifting the inhibitory / excitatory balance towards increased excitation via distinct mechanisms.

The other opioid receptor agonist investigated here was buprenorphine. This drug has a complex pharmacology, acting as a very high affinity partial agonist of the µ-OR, but an antagonist of δ- and κ-ORs (Davis, 2025). Buprenorphine injection caused a similar increase in beta-frequency LFP power to that observed after tianeptine, but with a peak effect at ∼50 min after administration. This is consistent with the relatively slow pharmacokinetics of buprenorphine which shows a half-life of several hours even after IV injection in rats (Gopal et al., 2002). Although buprenorphine’s antagonism of κ-ORs has been implicated in its antidepressant actions (e.g. Fava et al., 2016; but see Robinson et al., 2017), this action is unlikely to contribute to the increase in beta-frequency activity seen here since naloxone, another κ-OR antagonist (Lambert, 2023), had no effect on LFP power when administered alone or in combination with buprenorphine. Taken together, these results are however consistent with the idea that the µ-OR is the key target underlying buprenorphine and tianeptine’s LFP effects.

In contrast, the effects of ketamine on LFP and fEPSP measures were distinct from those of tianeptine and buprenorphine. Ketamine caused an increase in gamma power, consistent with previous studies of its hippocampal and cortical effects (e.g. Ma & Leung, 2007; Caixeta et al., 2013). There is evidence that ketamine’s antidepressant effects may be dependent on opioid receptor activation (Williams et al., 2018; Klein et al., 2020; Zhang et al., 2021; Adzic et al., 2023); both ketamine and its therapeutically active metabolite 2,6-hydroxynorketamine (2,6-HNK) are positive allosteric modulators of the µ-OR (Gomes et al., 2024), suggesting a role for the endogenous opioid system in ketamine’s antidepressant effects. With respect to its actions on LFP activity, however, ketamine’s enhancement of hippocampal gamma oscillations was here unaffected by a dose of naloxone that completely blocked the LFP changes associated with tianeptine and buprenorphine. Ketamine also caused a modest decrease in the fEPSP slope and amplitude. This may reflect a postsynaptic action at NMDA receptors located on CA1 pyramidal neurons. Previous studies have indicated a larger inhibitory effect of NMDA receptor blockade on baseline excitatory synaptic transmission in vivo, compared to in vitro (Bast et al., 2005). These fEPSP changes were also insensitive to naloxone in the current study.

A limitation of the present study is the use of urethane anaesthesia. Urethane has a mixed pharmacology, potentiating the activity of GABA_A_ receptors, and inhibiting AMPA and NMDA receptors (Hara & Harris, 2002). Under urethane, hippocampal LFP activity is characterised by a pattern of large-amplitude irregular activity (LIA) reminiscent of that observed during slow-wave sleep, interspersed with episodes of REM-sleep-like atropine-sensitive theta activity (Pagliardini et al., 2013). Relative to the awake state, spectral power is typically shifted towards lower frequencies under urethane anaesthesia; simplistically, this might reflect a prolongation of GABA_A_-receptor-mediated currents by urethane and a corresponding slowing of oscillatory network dynamics. Accordingly, our preliminary data suggest that tianeptine produces similar changes in LFP power in awake mice, but at the higher end of the beta-frequency range (Trigo et al., 2022); this study included both male and female animals, confirming that similar effects are seen in both sexes, as well as in both rodent species. One advantage of the use of anaesthesia in the current study is that the influence of locomotor activity on LFP activity can be definitively excluded (see Crouch et al., 2019). Since tianeptine is a locomotor stimulant—an effect abolished by knockdown of µ-opioid receptors on dopamine-D1-receptor-expressing medium spiny neurons of the striatum (Han et al., 2022)—this presents an additional complication in studies of awake animals.

The link between opioid receptor activation and network activity remains an underexplored area of research, but our current results provide clear evidence that the atypical antidepressant tianeptine causes beta-frequency oscillations in an opioid-receptor-dependent manner. An increased focus on the systems-level roles of hippocampal opioid receptors, or their downstream signalling pathways, might yield further insights into their participation in processes such as memory formation, and point towards new therapeutic strategies for the treatment of depressive illness.

## Acknowledgements

This work was supported by a Tenovus Scotland PhD studentship (T18/10) awarded to Stephen Martin and Jeremy Lambert and held by Scott Burt. We would like to thank the staff of the Medical Sciences Resource Unit for animal husbandry and welfare, Patrick Spooner for LabView programming, and Tim Hales for discussion of the project and comments on a draft of this manuscript.

## Data availability

Data are available on request.

