## Supplementary material for "The atypical antidepressant tianeptine causes opioid-receptor-dependent beta oscillations in the rat hippocampus"

### fEPSP measures of excitatory and inhibitory transmission

Fig. S1A shows a schematic illustration of the feedforward recruitment of inhibitory interneurons as well as excitatory neurotransmission after Schaffer collateral stimulation. A hypothetical example of the summation of the resulting IPSP and EPSP components, resulting in a dual-component fEPSP, is shown in Fig. S1B (cf. Alger & Nicoll, 1982; Arai et al., 1995; Karnup & Stelzer, 1999). The amplitude of the overshoot portion of the fEPSP tail provides a measure of the strength of feedforward inhibitory transmission.

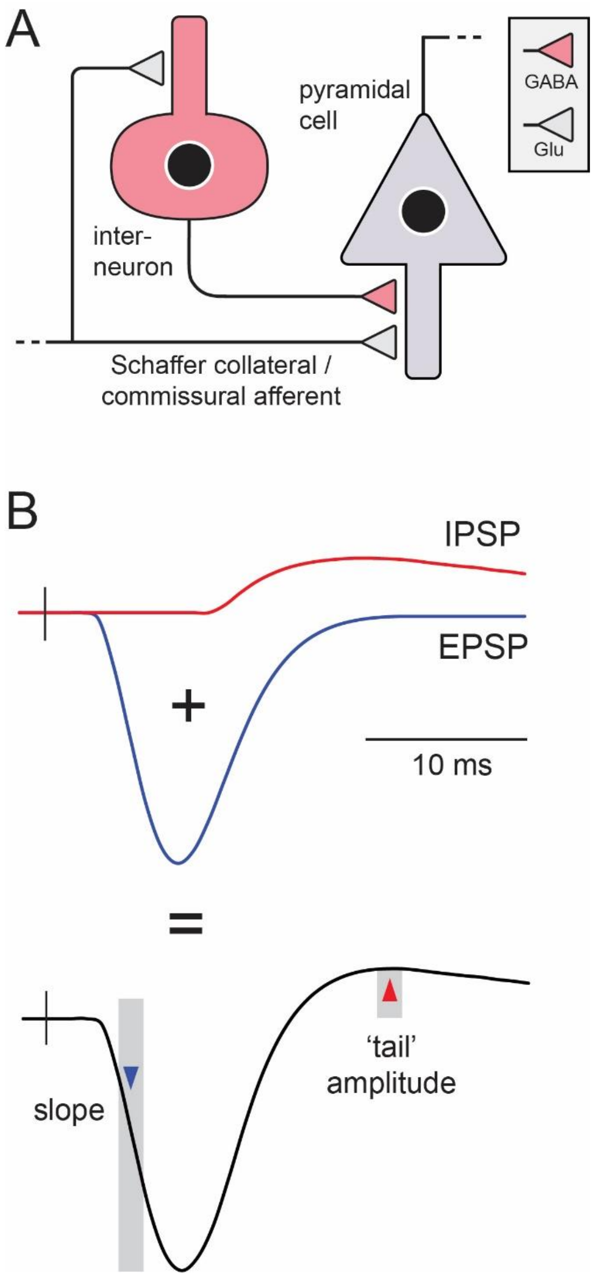


**Fig. S1.** Explanation of the dual-component fEPSP originating from Schaffer collateral stimulation. (A) Stimulation of Schaffer collaterals results in the activation of glutamatergic synapses located on CA1 pyramidal cells, but also the recruitment of inhibitory interneurons, leading to the feedforward activation of inhibitory inputs to the pyramidal cell. (B) The upper panel shows hypothetical extracellular EPSP and IPSP components recorded in the stratum radiatum of CA1 reflecting a negative-going monosynaptic excitatory EPSP (blue line) and a longer-latency positive-going inhibitory IPSP. The summation of these two components results in the compound EPSP shown in the lower panel. The slope of the early rising phase of the EPSP (blue triangle; measured by linear regression over the region indicated by the blue rectangle) was used as a measure of excitatory synaptic transmission in Fig. 4 & 7, and the amplitude of the positive-going EPSP tail (relative to the baseline preceding the stimulus artifact) was used as a measure of the strength of feedforward inhibition.

### Baseline values of electrophysiological measures

Table S1 shows baseline measures for the electrophysiological measures reported in the main text and supplementary material. Comparisons that reached significance by chance are indicated in red, and are further discussed below.

| | number | fEPSP slope (mV/ms) | fEPSP tail amplitude (AU) | theta ($log_{10}$ $\mu V^2/Hz$) | beta ($log_{10}$ $\mu V^2/Hz$) | slow gamma ($log_{10}$ $\mu V^2/Hz$) | fast gamma ($log_{10}$ $\mu V^2/Hz$) |
|---|---|---|---|---|---|---|---|
| vehicle | 12 | -0.87 ± 0.12 | 0.144 ± 0.049 | 3.18 ± 0.07 | 2.11 ± 0.07 | 1.73 ± 0.07 | 0.92 ± 0.05 |
| 10 mg/kg tianeptine | 10 | -0.83 ± 0.09 | 0.085 ± 0.009 | 3.15 ± 0.08 | 2.08 ± 0.05 | 1.76 ± 0.06 | 0.90 ± 0.04 |
| 30 mg/kg tianeptine | 10 | -1.05 ± 0.13 | 0.131 ± 0.015 | 3.21 ± 0.12 | 2.12 ± 0.05 | 1.78 ± 0.05 | 0.89 ± 0.05 |
| Effect of group | N/A | F(2,29) = 0.97; p = 0.39 | F(2,29) = 0.86; p = 0.43 | F(2,29) = 0.13; p = 0.88 | F(2,29) = 0.11; p = 0.05 | F(2,29) = 0.21; p = 0.82 | F(2,29) = 0.11; p = 0.05 |
| 30 mg/kg tianeptine | 10 | -1.05 ± 0.13 | 0.131 ± 0.015 | 3.21 ± 0.12 | 2.12 ± 0.05 | 1.78 ± 0.05 | 0.89 ± 0.05 |
| 30 mg/kg tianeptine + 1 mg/kg naloxone | 7 | -0.72 ± 0.11 | 0.105 ± 0.026 | 3.21 ± 0.11 | 2.19 ± 0.13 | 1.86 ± 0.12 | 0.90 ± 0.09 |
| naloxone only | 8 | -0.90 ± 0.15 | 0.119 ± 0.0197 | 3.20 ± 0.06 | 2.23 ± 0.07 | 1.92 ± 0.08 | 0.98 ± 0.06 |
| Effect of group | N/A | F(2,22) = 1.44; p = 0.26 | F(2,22) = 0.44; p = 0.65 | F(2,22) = 0.005; p = 1.0 | F(2,22) = 0.48; p = 0.62 | F(2,22) = 0.88; p = 0.43 | F(2,22) = 0.51; p = 0.61 |
| 0.03 mg/kg buprenorphine | 7 | -0.78 ± 0.66 | 0.095 ± 0.027 | 3.73 ± 0.12 | 2.26 ± 0.06 | 1.92 ± 0.04 | 1.04 ± 0.04 |
| 0.03 mg/kg buprenorphine + 1 mg/kg naloxone | 5 | -0.66 ± 0.15± | 0.102 ± 0.012 | 3.12 ± 0.20 | 2.03 ± 0.10 | 1.78 ± 0.09 | 0.85 ± 0.09 |
| Effect of group | N/A | t(10) = 0.59; p = 0.57 | t(10) = 0.22; p = 0.83 | t(10) = 2.78; p = 0.020 | t(10) = 2.02; p = 0.071 | t(10) = 1.74; p = 0.11 | t(10) = 2.26; p = 0.048 |
| 10 mg/kg ketamine | 12 | -1.03 ± 0.11 | 0.120 ± 0.030 | 3.11 ± 0.05 | 2.11 ± 0.09 | 1.75 ± 0.09 | 0.90 ± 0.07 |
| 10 mg/kg ketamine + 1 mg/kg naloxone | 7 | -1.01 ± 0.18 | 0.099 ± 0.018 | 3.28 ± 0.07 | 2.22 ± 0.06 | 1.88 ± 0.07 | 0.96 ± 0.05 |
| Effect of group | N/A | t(17) = 0.08; p = 0.94 | t(17) = 0.51; p = 0.62 | t(17) = 1.87; p = 0.079 | t(17) = 0.85; p = 0.41 | t(17) = 1.01; p = 0.33 | t(17) = 0.62; p = 0.54 |

**Table S1.** Baseline values for electrophysiological measures reported in this study. The fEPSP tail amplitude is expressed in arbitrary units (ratio of the positive tail amplitude and the negative phase of the fEPSP). Significant values are indicated in red.

### Changes in theta (3-4 Hz) and fast gamma (50-80 Hz) power

The analysis presented in the main text (Fig. 2, 3, 5 & 6) focusses on the beta and slow-gamma ranges since the most prominent changes in power were observed within these ranges. For completeness, we present an analysis of theta and fast-gamma changes below.

Fig. S2A shows the change in theta-frequency power after injection of vehicle or tianeptine at doses of 10 mg/kg and 30 mg/kg. The frequency range analysed (3-4 Hz) is indicated in the inset. All groups showed a transient increase in theta power after injection typically lasting several minutes and likely reflecting the activation of ascending nociceptive pathways by subcutaneous injection. However, an analysis of the mean change in theta power 20-30 min after injection revealed no overall group difference [$F(2,29) = 0.22$; $p = 0.81$], and one-sample t-tests with Bonferroni correction did not reveal any changes in power relative to baseline levels in any of the groups. Fig S2B shows a corresponding analysis of fast gamma power. Tianeptine caused a dose-dependent increase in fast gamma power, indicated by a significant main effect of tianeptine dose 20-30 min after injection [$F(2,29) = 17.0$; $p < 0.0001$], and post-hoc pairwise comparisons (Tukey's HSD) revealed significant differences between all groups (vehicle versus 10 mg/kg tianeptine: $p = 0.023$; vehicle versus 30 mg/kg tianeptine: $p < 0.0001$; 10 mg/kg tianeptine versus 30 mg/kg tianeptine: $p = 0.019$). There was a significant increase in fast gamma power relative to baseline in the 30 mg/kg tianeptine group [$t(9) = 5.55$; $p = 0.0012$; one-sample t-test with Bonferroni correction].

Fig. S2C shows the changes in theta power in rats injected with 30 mg/kg tianeptine + 1 mg/kg naloxone, or 1 mg/kg naloxone only. The 30 mg/kg tianeptine group from panel A is included for comparison. An ANOVA of mean theta power 20-30 min after injection in these 3 groups did not reveal a significant group effect [$F(2,22) = 2.79$; $p = 0.083$], and one-sample t-tests with Bonferroni correction did not reveal any changes in power relative to baseline levels in any of the groups. Fig S2D shows the changes in fast gamma power in rats injected with 30 mg/kg tianeptine + 1 mg/kg naloxone, or 1 mg/kg naloxone only. The 30 mg/kg tianeptine group from panel B is included for comparison. Co-administration of naloxone completely blocked the tianeptine-induced increase in fast gamma power. An ANOVA of the mean power 20-30 min after injection in the 3 groups revealed a significant effect of group [$F(2,22) = 15.4$; $p < 0.0001$], and significantly lower power in the tianeptine + naloxone ($p = 0.0028$) and naloxone only groups ($p = 0.00041$), neither of which showed a significant change in power relative to baseline (one-sample t-tests with Bonferroni correction).

Fig. S2E shows the mean change in theta power after injection of 0.03 mg/kg buprenorphine or 0.03 mg/kg buprenorphine + 1 mg/kg naloxone. Analysis of the mean change in theta power 30-60 min after injection revealed a significant reduction in power in the buprenorphine relative to the buprenorphine + naloxone group [$t(10) = 2.93$; $p = 0.015$], though the change relative to baseline values did not reach significance in either group (one-sample t-tests with Bonferroni correction). However, a complication is that baseline theta power was higher, by chance, in the buprenorphine group relative to the buprenorphine + naloxone group [$3.74 \pm 0.12$ versus $3.12 \pm 0.20$ $\log_{10}\mu V^2/Hz$; $t(10) = 2.78$; $p = 0.020$; see table S1]. This difference can be seen in Fig. 5A(ii) as a small peak at ~4 Hz in the power spectrum before buprenorphine injection, a peak that is less prominent in the buprenorphine + naloxone group [Fig. 5B(ii)]. The change in theta power was strongly negatively correlated with baseline theta power in both the 0.03 mg/kg buprenorphine [$r = -0.96$; $n = 7$; $p = 0.00075$] and 30 mg/kg tianeptine groups [$r = -0.78$; $n = 10$; $p = 0.0077$], although baseline theta power was significantly lower in the former case [$3.21 \pm 0.12$ versus $3.74 \pm 0.12$ $\log_{10}\mu V^2/Hz$ in the buprenorphine group; $t(15) = 2.95$; $p = 0.01$]. Because of this, the apparent difference between the effects of buprenorphine and tianeptine on theta power (Fig. S2A & E) might be a consequence of the higher baseline theta power in the buprenorphine group. In fact, splitting the 30 mg/kg tianeptine and 0.03 mg/kg buprenorphine groups into sub-groups with baseline theta power greater than the median in the former case and less than the median in the latter case yielded groups with equivalent baseline theta power [tianeptine: $3.50 \pm 0.14$ $\log_{10}\mu V^2/Hz$; $n = 5$; buprenorphine: $3.52 \pm 0.07$ $\log_{10}\mu V^2/Hz$; $n = 4$; $t(7) = 0.16$; $p = 0.87$]; these two sub-groups showed an equivalent change in theta power after injection [tianeptine: $-0.16 \pm 0.15$ $\log_{10}\mu V^2/Hz$; $n = 5$; buprenorphine: $-0.20 \pm 0.13$ $\log_{10}\mu V^2/Hz$; $n = 4$; $t(7) = 0.24$; $p = 0.82$]. However, the experimental procedures used here are not optimal for the assessment of theta-frequency changes owing to the sporadic nature of the theta state under urethane anaesthesia; the issue could be overcome by the induction of theta via brainstem stimulation, or the use of awake animals undergoing forced locomotion.

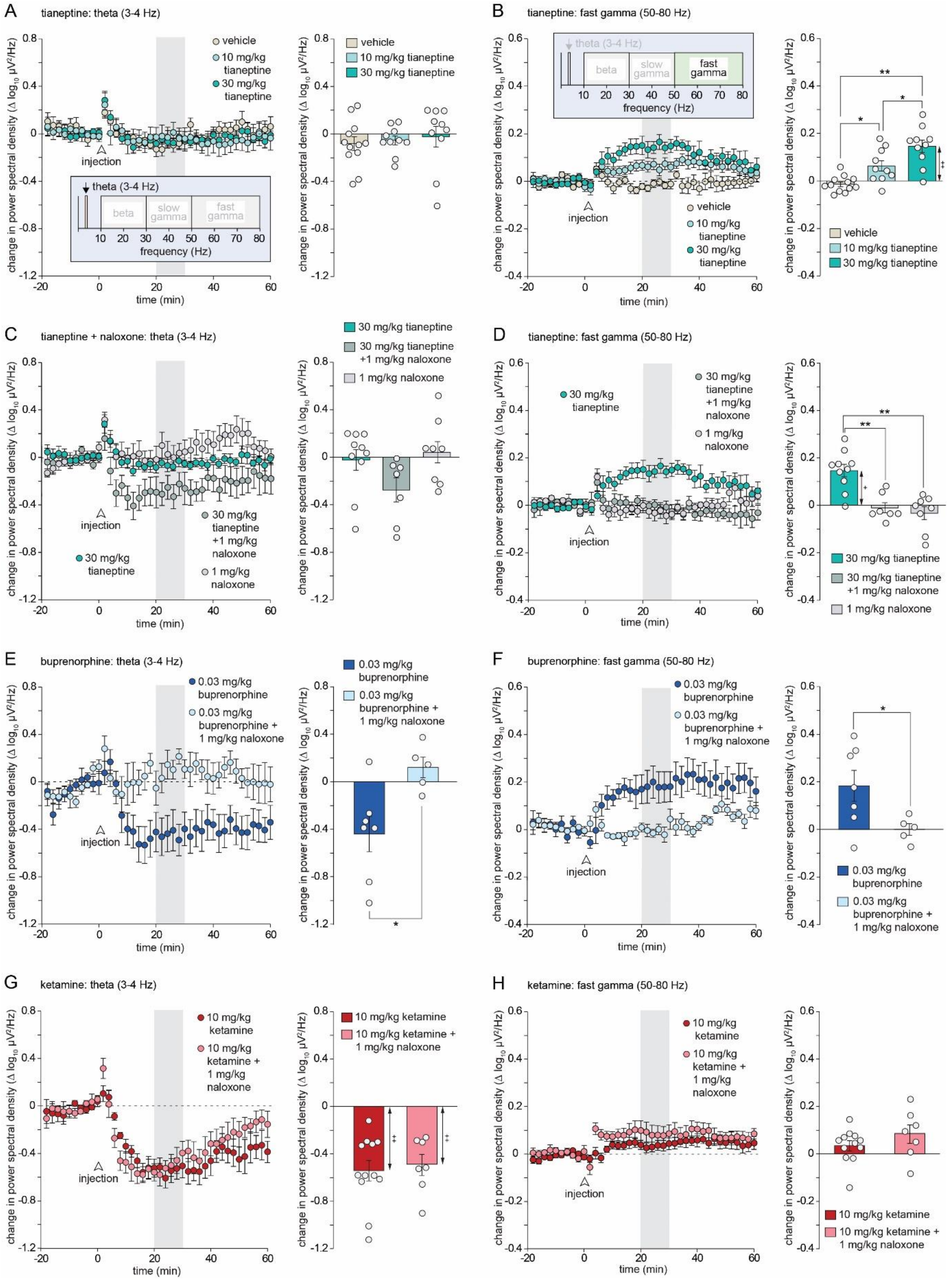


**Fig. S2.** Changes in theta (3-4 Hz) and fast gamma (50-80 Hz) oscillations. (A) Left-hand panel: time-course of changes in theta-frequency power (see inset) after administration of saline (n = 12), 10 mg/kg tianeptine (n = 10) and 30 mg/kg tianeptine (n = 10); right-hand panel: mean change in theta power 20-30 min (corresponding to the grey bar in the left-hand panel) after injection relative to baseline. (B) Left-hand panel: time-course of changes in fast gamma power (see inset) after administration of the same drugs as in A; right-hand panel: mean change in slow gamma power 20-30 min after injection relative to baseline.

(C) Left-hand panel: time course of changes in theta power after injection of 30 mg/kg tianeptine + 1 mg/kg naloxone ($n = 7$), naloxone only ($n = 8$), or 30 mg/kg tianeptine ($n = 10$; data reproduced from A for comparison); right-hand panel: mean change in theta power 20-30 min after injection relative to baseline. (D) Left-hand panel: time course of changes in fast gamma power after injection of the same drugs as in C; right-hand panel: mean change in fast gamma power 20-30 min after injection relative to baseline. (E) Left-hand panel: time-course of changes in theta power after injection of 0.03 mg/kg buprenorphine ($n = 7$) or 0.03 mg/kg buprenorphine + 1 mg/kg naloxone ($n = 5$); right-hand panel: mean change in theta power 20-30 min after injection relative to baseline. (F) Left-hand panel: time-course of changes in fast gamma power after injection of the same drugs as in E; right-hand panel: mean change in theta power 20-30 min after injection relative to baseline. (G) Time-course of changes in theta power after injection of 10 mg/kg ketamine ($n = 12$) or 10 mg/kg ketamine + 1 mg/kg naloxone ($n = 7$); right-hand panel: mean change in theta power 20-30 min after injection relative to baseline. (H) Left-hand panel: time-course of changes in fast gamma after injection of the same drugs as in G; right-hand panel: mean change in theta power 20-30 min after injection relative to baseline. *$p < 0.05$; **$p < 0.01$: independent samples t-test or post-hoc pairwise comparisons (Tukey's HSD); +$p < 0.05$; ++$p < 0.01$: one-sample t-test with Bonferroni correction for multiple comparisons.

Fig. S2F shows the mean change in fast gamma power after injection of 0.03 mg/kg buprenorphine or 0.03 mg/kg buprenorphine + 1 mg/kg naloxone. Analysis of mean power 20-30 min after injection revealed that naloxone co-administration caused a significant reduction in power relative to the buprenorphine only group [$t(10) = 2.29$; $p = 0.045$]. However the change in power relative to baseline did not reach significance in either group. Although there was a significant difference in baseline fast-gamma power between the buprenorphine and buprenorphine + naloxone groups ($3.73 \pm 0.12$ $\log_{10}\mu V^2/Hz$ and $3.12 \pm 0.20$ $\log_{10}\mu V^2/Hz$ respectively; $t(10) = 2.26$; $p = 0.048$; see table S1), it is unlikely that this can account for naloxone's reduction in fast gamma activity because there was no significant relationship between baseline fast gamma power in the buprenorphine group and the change in power after injection ($r = 0.0063$; $n = 7$; $p = 0.99$).

Fig. S2G shows the mean change in theta power after injection of 10 mg/kg ketamine or co-injection of 10 mg/kg ketamine + 1 mg/kg naloxone. Ketamine caused a decrease in theta-frequency power that was not blocked by naloxone. Both groups showed a significant decrease in mean power 20-30 min after injection (ketamine: $p = 0.0002$; ketamine + naloxone: $p = 0.0028$; one-sample t-tests with Bonferroni correction), and there was no significant difference between them [$t(17) = 0.37$; $p = 0.71$]. Fig. 2SH shows the corresponding changes in fast gamma power for these groups. Neither group showed a significant change in fast gamma power relative to baseline values (one-sample t-tests with Bonferroni correction), and there was no difference between them [$t(17) = 1.48$; $p = 0.15$].

**Relationship between brain temperature and fEPSP measures (slope and tail amplitude)**

In the subset of vehicle-treated rats implanted with temperature probes, once the main experiment was completed, the heat blanket was used to passively warm the animal for a period of 14-18 min, typically sufficient to cause an increase of ~0.8°C, i.e. within the range typically caused by tianeptine administration. Stimulation pulses were delivered every 20 s with parameters identical to those used in the main experiment. The fEPSP slope was normalised to the value over the first 1 min of the recording period and plotted against the mean increase in brain temperature, also normalised to the first 1 min of recording, and averaged over the 20 s after each stimulation pulse.

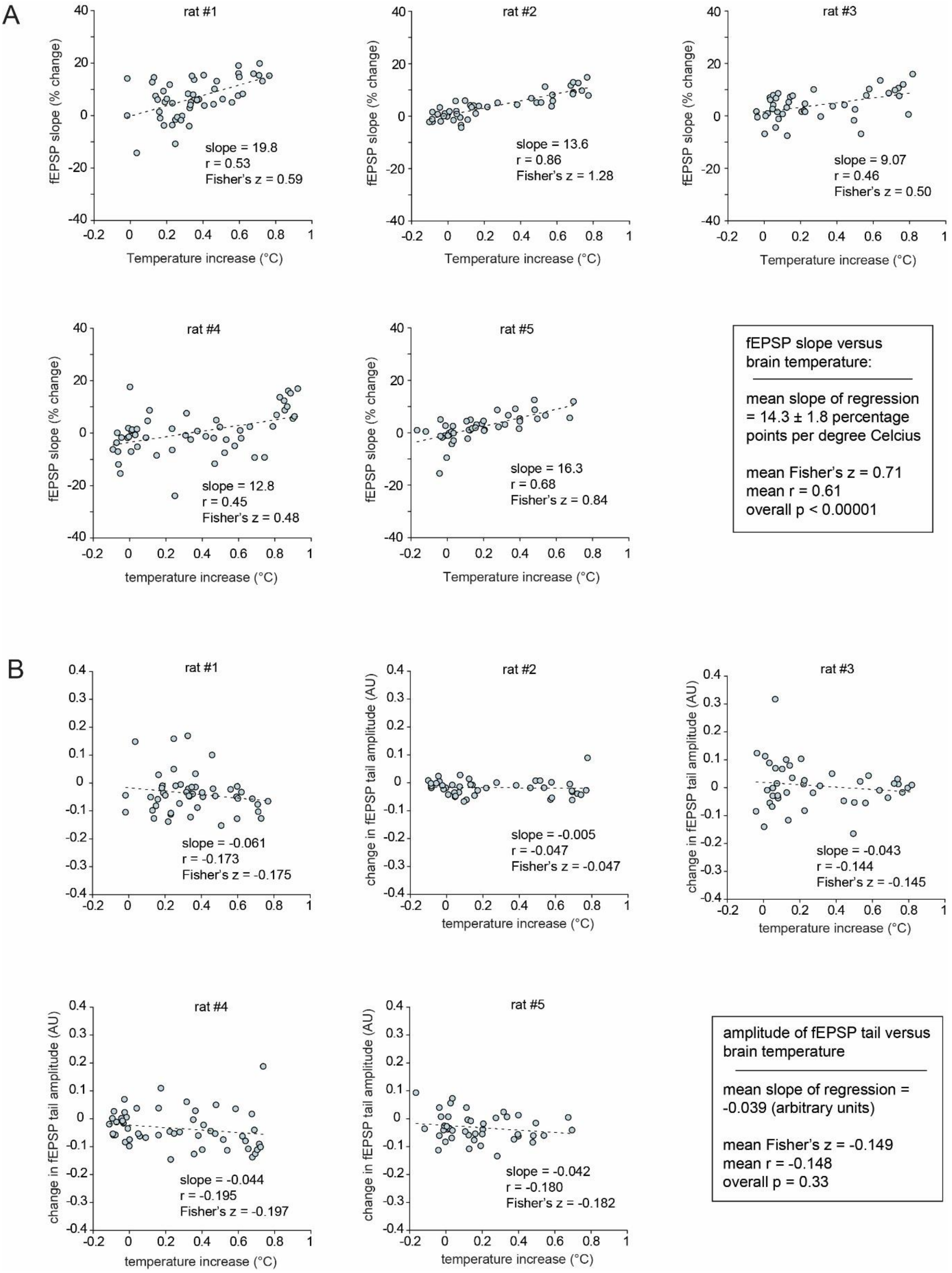


**Fig. S3.** Relationship between brain temperature and LFP activity. (A) Positive correlation between the increase in fEPSP slope and brain temperature during passive warming in 5 individual rats. The overall Peason's correlation coefficient was highly significant. (B) Corresponding correlation between fEPSP tail amplitude and brain temperature. The weak negative correlation was not statistically significant.

Fig. S3A shows the correlations between fEPSP slope and brain temperature in each rat. The mean slope of the relationship was 14.10 ± 1.89 percentage points per degree of temperature increase, a value significantly above zero [$t(4) = 7.46$; $p = 0.0017$; one-sample t-test]. To calculate a mean correlation coefficient, individual r values were converted to Fisher's z values, a weighted mean was calculated based on the number of samples, and this mean was converted back to an overall r value. A highly significant positive correlation between fEPSP slope and brain temperature was observed [mean Fisher's $z = 0.71$; mean $r = 0.61$; $n = 46$; $p < 0.00001$]. An equivalent analysis of the relationship between fEPSP tail amplitude and brain temperature is shown in Fig. S3B. There was a weakly negative, but non-significant, relationship between these measures [mean Fisher's $z = -0.149$; mean $r = -0.148$; $n = 46$; $p = 0.33$].

Based on the mean slope of the relationship between fEPSP slope and brain temperature, a predicted increase in fEPSP was calculated for the rats in which brain temperature was recorded during injection of tianeptine and buprenorphine. Table S2A shows the mean observed and predicted values 20-30 min after injection. These values did not differ in any group, indicating that the small increases in fEPSP observed after tianeptine and buprenorphine injection are entirely accounted for by the increase in brain temperature. In contrast, the decrease in tail amplitude in the 30 mg/kg tianeptine group was significantly larger than predicted based on the temperature increase, and no-significant trends in this direction were observed in the 10 mg/kg and buprenorphine groups (Table S2B). This suggests that while the increase in fEPSP slope is caused entirely by the increase in brain temperature, separate mechanisms underlie the tianeptine-induced reduction in tail amplitude, such as an inhibition of feedforward inhibition.

| A: fEPSP slope | 10 mg/kg tianeptine | 30 mg/kg tianeptine | 0.03 mg/kg buprenorphine |
|---|---|---|---|
| observed temperature increase | 0.14 ± 0.09 °C | 0.56 ± 0.13 °C | 0.45 ± 0.05 °C |
| observed fEPSP slope (% baseline) | 1.44 ± 2.37% | 7.89 ± 2.38% | 10.36 ± 5.60% |
| predicted fEPSP slope (% baseline) | 2.01 ± 1.28% | 7.99 ± 1.88% | 6.02 ± 0.80% |
| t value (paired-sample) | 0.23 | 0.0032 | 0.68 |
| sample size | 6 | 6 | 7 |
| p value | 0.83 | 0.98 | 0.52 |

| B: fEPSP tail amplitude | 10 mg/kg tianeptine | 30 mg/kg tianeptine | 0.03 mg/kg buprenorphine |
|---|---|---|---|
| observed temperature increase | 0.14 ± 0.09 °C | 0.56 ± 0.13 °C | 0.45 ± 0.05 °C |
| observed change in tail amplitude (AU) | -0.027 ± 0.009% | -0.067 ± 0.019% | -0.072 ± 0.025% |
| predicted change in tail amplitude (AU) | -0.005 ± 0.003% | -0.022 ± 0.005% | -0.016 ± 0.002% |
| t value (paired-sample) | 2.52 | 2.58 | 2.06 |
| sample size | 6 | 6 | 7 |
| p value | 0.053 | 0.049 | 0.085 |

**Table S2.** Observed and predicted mean fEPSP slope (A) and tail amplitude values (B) 20-30 min after drug injection in the subsets of tianeptine- and buprenorphine-treated rats implanted with a temperature probe. Predicted values are based on the observed temperature increase as explained in the text above. Red text indicates a statistically significant effect.

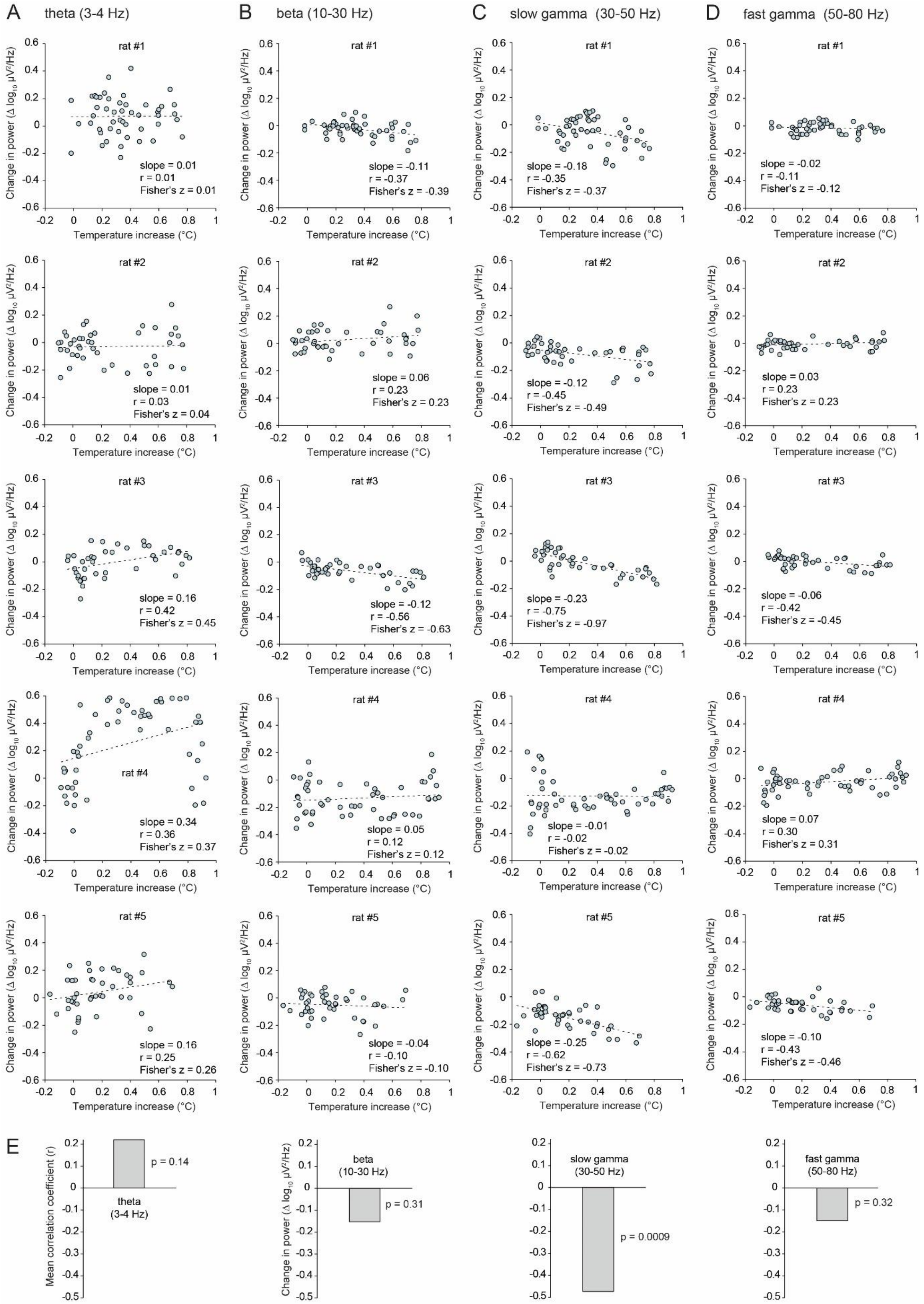


**Fig. S4.** Relationship between brain temperature and LFP power. Correlations between brain temperature during passive warming in 5 individual rats are shown for the theta (A), beta (B), slow gamma (C), and fast gamma (D) ranges. (E) Histogram of the overall value of Pearson's correlation coefficient (r); a significant negative correlation between brain temperature and LFP power was evident in the slow-gamma band.

Fig. S4 shows the correlations between the increase in log power in each LFP frequency band and brain temperature. Statistical analysis was carried out in the same way as described above for fEPSP slope. There was a weak positive correlation between brain temperature and theta power (Fig. S4A), and the slope of the correlation was negative in the beta (Fig. S4B), slow gamma (Fig. S4C) and fast gamma (Fig. S4D) ranges. However, a significant association was only observed in the slow-gamma band. The mean slope of the regression line was -0.16 ± 0.04 power spectral density units ($log_{10}$ $\mu V^2/Hz$) per degree of temperature increase, a value significantly below zero [$t(4) = 3.60$; $p = 0.023$; one-sample t-test]. The overall correlation coefficient derived from the mean Fisher's z score was highly significant [mean Fisher's $z = -0.52$; mean $r = -0.47$; $n = 46$; $p = 0.00088$]. As shown in Fig. S4, the relationship between spectral power and brain temperature did not reach significance in other frequency bands.

Fig. S4D summarises the strength of the relationship between brain temperature and LFP power across different frequency bands by plotting the mean Pearson's r value for each band. This analysis reveal a U-shaped relationship between brain temperature and change in power across the spectral frequency range. Note that the negative relationships between LFP power and temperature in the beta and gamma frequency ranges predict a tianeptine-induced fall in spectral power based on drug-induced temperature increases alone. This contrasts sharply with the observation that tianeptine causes a pronounced in beta-frequency power.
